# Contribution of cytoplasm viscoelastic properties to mitotic spindle positioning

**DOI:** 10.1101/2021.10.21.465315

**Authors:** Jing Xie, Javad Najafi, Rémi Le Borgne, Jean-Marc Verbavatz, Catherine Durieu, Jeremy Sallé, Nicolas Minc

## Abstract

Cells are filled with macromolecules and polymer networks that set scale-dependent viscous and elastic properties to the cytoplasm. Although the role of these parameters in molecular diffusion, reaction kinetics and cellular biochemistry is being increasingly recognized, their contributions to the motion and positioning of larger organelles, such as mitotic spindles for cell division remain unknown. Here, using magnetic tweezers to displace and rotate mitotic spindles in living embryos, we uncovered that the cytoplasm can impart viscoelastic reactive forces that move spindles, or passive objects with similar size, back to their original position. These forces are independent of cytoskeletal force generators, yet reach hundreds of piconewtons and scale with cytoplasm crowding. Spindle motion shears and fluidizes the cytoplasm, dissipating elastic energy and limiting spindle recoils with functional implications for asymmetric and oriented divisions. These findings suggest that bulk cytoplasm material properties may constitute important control elements for the regulation of division positioning and cellular organization.

**Significance Statement:** The regulation of mitotic spindle positioning is a key process for tissue architecture, embryo development and stem cells. To date, most models have assumed that spindles are positioned by forces exerted by polar cytoskeleton networks, like microtubule asters or acto-myosin bundles. Here, using *in situ* magnetic tweezers to apply calibrated forces and torques to mitotic spindles in live dividing sea urchin cells, we found that the viscoelastic properties of the cytoplasm medium in which spindles are embedded can hold spindles in place, and move them back if their original position is perturbed. These viscoelastic forces are large and may significantly participate in the force balance that position and orient mitotic spindles in many cell types.

## Introduction

The cytoplasm is a heterogeneous composite material crowded with large macromolecular complexes, endomembranes, and entangled cytoskeletal networks (1, 2). These set a hierarchy of pore and mesh sizes which define rheological properties, such as viscosity and elasticity, that impact fundamental processes ranging from the kinetics of biochemical reactions to vesicular transport and cell shape control (3-5). The importance of cytoplasm material properties for cellular physiology has been recognized and studied for decades, starting from early micro-rheology experiments by Crick or Hiramoto (6-8). These showed that injected micrometric beads displaced in the cytoplasm exhibit typical viscoelastic responses, with partial positional feedback that move them back towards their initial position. Thus, the cytoplasm features both solid-like and fluid-like behavior, with bulk elastic moduli on the order of ∼1-10 Pa, typical of soft gels and viscosities 100-1000 times that of water. More recent studies have now established that these rheological characteristics exhibit size, force or frequency dependence, and provided more refined descriptions of the cytoplasm using frameworks of non-linear viscoelasticity or poroelasticity (2, 3, 9-11). Object size is of particular relevance, given that components floating in the cytoplasm may range over 4-5 orders of magnitudes. Indeed, cytoplasm rheology has been proposed to transit from that of a Newtonian fluid for small particles to that of a more glassy or elastic solid for larger elements (11, 12). To date, however, many studies of bulk cytoplasm properties and their functions have focused on relatively small objects, leaving the fundamental problem of how they impact the motion of large organelles, like nuclei or cytoskeletal assemblies poorly explored.

The mitotic spindle is one such large assembly that resides at a precise location in the cytoplasm to specify cytokinesis, and thus the size and position of daughter cells in tissues (13, 14). Spindles are built from dynamic microtubules (MTs) and motors and can take up significant portions of cellular space. They are commonly associated with networks of nuclear intermediate filaments and endomembranes that form a so-called spindle matrix (15, 16). These considerations suggest that their motion in the dense cytoplasm could be associated with large viscous and elastic drags, with potential implications for division positioning and chromosome segregation. Until now however, the literature covering the mechanics of spindle positioning has been dominated by the role of active directed forces from polar cytoskeletal networks (13, 14, 17). Spindles may for instance decenter or rotate, during asymmetric or oriented divisions, a process typically associated with forces generated by contractile acto-myosin networks (18, 19) or astral microtubules (MTs) and associated motors like dynein (20). For symmetric divisions, mitotic spindles reside stably in the cell center. This is thought to be regulated by MTs that grow to contact cell boundaries and exert length-dependent pushing and/or pulling forces on the spindle: when spindles become off-centered, asymmetries in MT lengths and forces act as an effective spring related to cell shape to re-center spindles (21-25). Net forces applied by MT asters or acto-myosin networks may range from few tens to hundreds of piconewtons (pN) (26-28). A displacement of an impermeable objects with the radius of a spindle of R= 5 µm by a distance d=5 µm, typical of many asymmetric divisions, in a medium with an elastic modulus of G=1 Pa, like the cytoplasm, would generate a reactive force F=6πR*G*d ∼500 pN. Thus, viscoelastic properties of the cytoplasm could in principle be highly relevant to the mechanics of spindle positioning. To date, however, the lack of proper assays to probe cytoplasm rheology at the scale of a moving spindle has impaired testing this fundamental problem for cell organization.

Here, by exploiting large sea urchin cells, where mitotic asters are too short to reach the cell surface, we establish and quantify the direct contribution of bulk cytoplasm viscoelasticity to the mechanics of spindle positioning. We use spindles or large passive oil droplets moved and rotated by calibrated magnetic tweezers in intact cells to probe cytoplasm viscosity and elasticity, at time and length scales representative of spindle movements commonly observed in asymmetric or oriented divisions. We find that the stress exerted by the spindle on the cytoplasm causes it to flow and deform, and exert large reactive spring-like forces that move back this large organelle towards its initial position. Cellular-scale flows also shear and rearrange the cytoplasm, dissipating elastic energy and rendering spindle repositioning time-dependent which facilitates rotational over translational spindle motions. Our results place cytoplasm rheology as a hitherto unappreciated element in the force balance that controls the positioning of mitotic spindle and potentially other large organelles.

## Results

### Viscoelastic forces maintain metaphase spindle position even in the absence of astral MTs contacting the cortex

In many small cells, mitotic spindles are connected to the cell cortex by dynamic MTs which act as dominant force generators to maintain or modulate spindle position (29). In larger cells, limits in spindle size may prevent bounded metaphase mitotic asters from reaching the cell surface (30-34). Using immunofluorescence and Airy-scan confocal microscopy to detect individual astral MTs around metaphase spindles of 95 µm-sized sea urchin zygotes, we computed a mean distance from astral MTs +tips to the actin-rich cortex of 14.5 +/-9.8 µm (+/-SD), which corresponds to ∼15 % of egg size. Out of ∼4000 MTs tracked we found a mean of only 3.35 +/-3.4 MTs/cell that came within 5 µm of the egg surface, a distance typically larger than the actin cortex in these eggs (Fig 1A-1B) (35). These results were confirmed by visualizing MTs in live cells with different probes, as well as with transmission electron microscopy of eggs fixed with optimized methods to reveal MTs (Fig S1A-S1D) (36). In sharp contrast, and as previously reported, interphase and anaphase/telophase asters spanned the whole cell with a mean of 373 +/-12 and 408 +/-33 MTs/cell reaching a distance less than 5 µm to the surface, respectively (Fig 1C and Fig S1E) (30, 33). In spite of lacking MT contact with the cell surface, the spindle appeared largely static at the cell center in both position and orientation, over the typical ∼10 min duration of metaphase, or over longer time-scales up to 35-40 min when metaphase was prolonged with the proteasome inhibitor MG132 (Movie S1). Thus, metaphase spindles can robustly maintain their position and orientation for long periods of times, even in the absence of astral MTs contacting the cell cortex.

**Figure 1.**
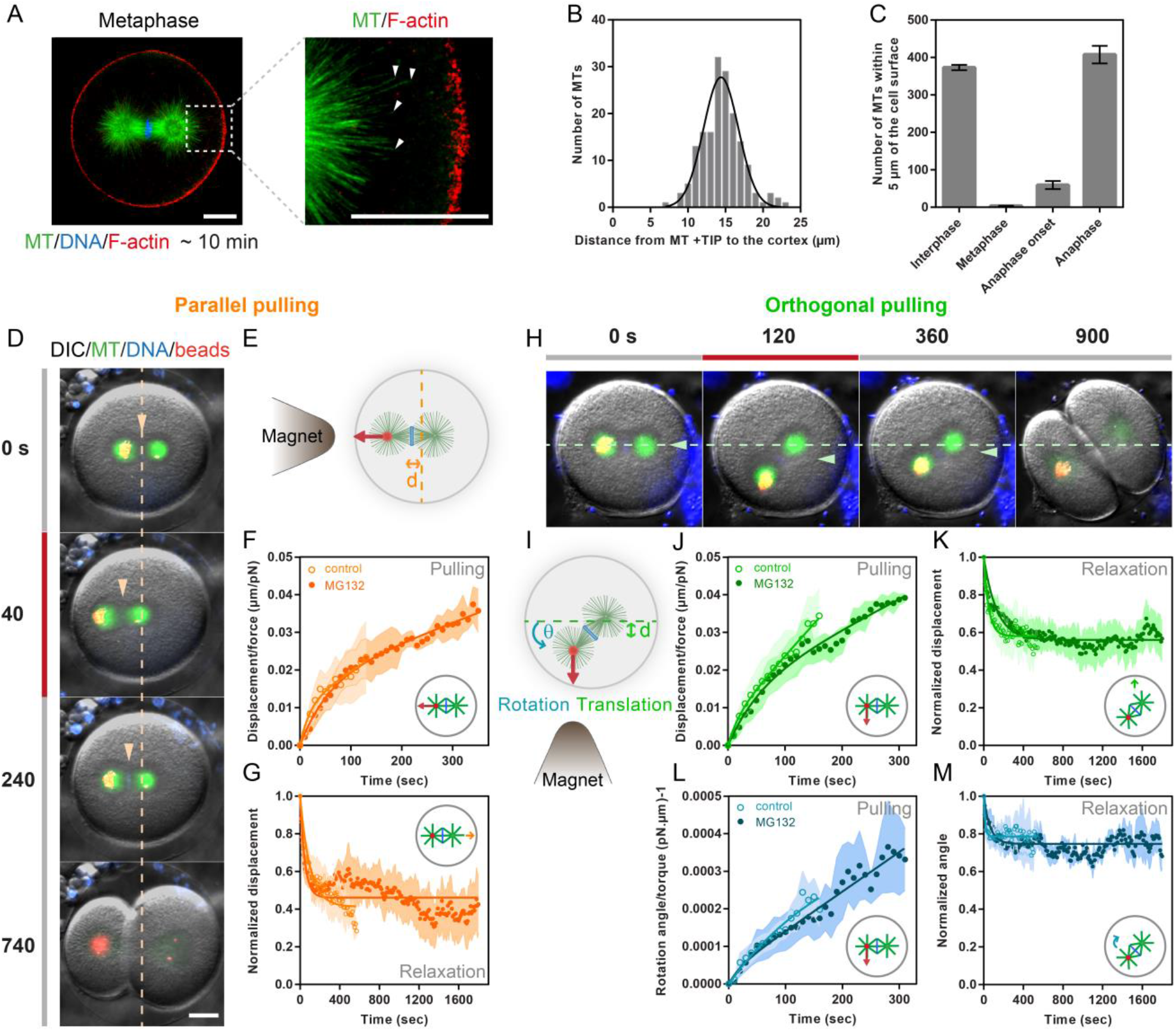
Viscoelastic forces hold spindles in the center of large cells even in the absence of astral Microtubules contacting the cell cortex. **(A)** (Left) Airy-scan confocal image of a sea urchin zygote in metaphase fixed and stained for Microtubules (MTs), DNA and F-actin. (Right) Close up view on + Ends of microtubules, marked with white arrowheads, and the actin-rich cortex. **(B)** Quantification of the distance from MT+ TIPS to the actin cortex (n= 168 MTs from 4 eggs). **(C)** Number of MTs reaching a distance less than 5 µm from the cortex in different phase of the first cell cycle (n= 3, 20, 5 and 2 cells respectively). Error bars correspond to +/-SEM. **(D-E)** Time-lapse of metaphase spindles with magnetic beads bound to one spindle pole, displaced by magnetic forces applied parallel to the spindle axis by the presence of a magnet tip, and recoiling upon force cessation. **(F)** Time evolution of the displacement measured from the initial centered position of the spindle normalized by the applied force for metaphase spindles in normal cells (n=11) and in cells treated with MG132 to arrest cells in metaphase (n=9). **(G)** Time evolution of the displacement back to the cell center when the external force is released, normalized to that at the moment of force cessation, for the same cells and conditions as in F. **(H-I)** Time-lapse of metaphase spindles displaced and rotated by magnetic forces applied orthogonal to the spindle axis, and spindle recoiling upon force cessation. **(J)** Time evolution of the displacement measured from the initial centered position of the spindle and normalized by the applied force in normal cells (n=14) and in cells treated with MG132 (n=7). **(K)** Time evolution of the normalized displacement back to the cell center when the external force is released for the same cells and conditions as in J. **(L)** Time evolution of the spindle axis angle normalized by the external torque applied in normal cells (n=14) and in cells treated with MG132 (n=7). **(M)** Time evolution of the normalized angle when the external torque is released for the same cells and conditions as in L. In F-G, and J-M, the bold lines correspond to fits of the data using general creep or relaxation equations of the Jeffreys’ viscoelastic model (see main text and methods). Error bars are represented as shades in these curves and correspond to +/-S.D/2. Scale bars, 20 µm.

To directly modulate spindle position and orientation, we implemented *in vivo* magnetic tweezers to apply forces and torques to spindles in live cells (26). We injected a specific type of magnetic beads in unfertilized eggs, and added sperm to trigger fertilization (28, 37). These beads exhibit spontaneous centripetal motion along MT asters and form compact aggregates that stay attached to centrosomes through the cell cycle in these cells, allowing to apply magnetic forces on centrosomes by approaching a magnet tip. These beads also form aggregates in vitro, which allows for the calibration of the magnetic force as function of aggregate size and distance to the magnet tip, by tracking beads velocities in test viscous fluids (37). The presence of beads at spindle poles did not affect spindle dimensions, and had no notable effect on cell cycle progression (Fig S1F-S1G). In some embryos, beads often split into two aggregates, following centrosome duplication in interphase or early prophase. In others, the beads only tracked one centrosome, allowing a point force application at a single spindle pole (Fig S1H-S1I). Using those, we applied external forces ranging from ∼70 to 700 pN along different axis, and monitored resultant spindle motion. Pulling the spindle parallel to its long axis, caused the spindle to translate towards the magnet tip, while a force applied orthogonal to the spindle axis caused both translational and rotational motions that tended to align the spindle along the magnetic force axis (Fig 1D-1E and 1H-1I, Movie S2 and S3). Therefore, these experiments allowed to recapitulate spindle movements typically observed in asymmetric or oriented divisions with calibrated forces and torques in intact cells.

Astral MTs that grow to the cortex, to push or pull on spindles, may act effectively as an active elastic system related to cell shape that brings back a spindle to the cell center if its position is perturbed (21, 23). In our system, where spindles lack MTs reaching cell boundaries, we anticipated a viscous response to applied forces with no elastic positional feedback. To test this, we collapsed displacement-time curves from individual spindle pulls under different force magnitudes, by rescaling spindle displacement by force (26). This rescaling also allowed to compensate for small variations (∼10-20%) in external forces during each pull. Strikingly, these rescaled displacement-time curves of spindles moved parallel or orthogonal to their long axis, exhibited a typical viscoelastic response: spindle motion was first linear at short time-scales below 10-30s, following a viscous regime, with an initial speed proportional to the applied force, but then slowed down, yielding an inflection in the displacement-time curve indicative of internal elastic forces that push or pull back the spindle to oppose external forces (Fig 1F, 1J and S1J). Accordingly, larger forces yielded larger displacements at a fixed time point in the inflecting regime, and when the force was released, the spindle recoiled back (Fig 1G, 1K and Fig S1K). We also noted that at longer time scales above ∼100-200 s the curve tended to converge onto another linear regime. In addition, recoils were only partial, with spindles recovering ∼ 40-60 % of their initial displacements, often yielding a small asymmetry in division plane positioning (Fig 1G and 1K). These behaviors reflect significant dissipations in the stored elastic energy.

Rotational dynamics of spindles submitted to magnetic torques also exhibited a viscoelastic response, but elastic recoils appeared less pronounced than in translation, causing spindles to tilt and mostly maintain their final orientation at the time of force release (Fig 1L and 1M). Importantly, similar responses were obtained in cells arrested in metaphase with MG132, ruling out putative contributions of aster regrowth and initial cortex contact in late metaphase. Spindle pulling assays were also limited to a small enough displacement that ensured that mitotic asters did not contact the cortex, and spindle re-centering did not exhibit any correlation with the final distance to the cortex (Fig 1F-1G, 1J-1M and Fig S1L). Thus, although these data cannot firmly reject a minor role of MTs contacting the cortex, they suggest that most of this viscoelastic response may be attributed to elements in the cytoplasm. Together these results suggest the existence of viscoelastic restoring forces that maintain spindle positon and orientation, even in the absence of MTs reaching the cell cortex.

### Spindle repositioning is caused by viscoelastic restoring forces from bulk cytoplasm material

To understand the origin of these viscoelastic restoring forces, we tested the role of MTs as prominent force-generators for spindle positioning. We displaced spindles with magnetic tweezers, and rapidly rinsed cells with Nocodazole, to affect MTs, and monitored the ability of spindles to recoil back (Fig 2A). In controls, the positional recovery followed a single exponential with a decay time-scale of 103 +/-92 s and a positional offset of 43 +/-24%. Nocodazole treated spindles shrank in size to eventually disassemble, over a period of ∼5-10 min, but recovered their positions with similar dynamics and offsets as controls (Fig 2B-2D and Fig S2A-S2E). Therefore, in agreement with the lack of MTs reaching cell boundaries, astral MT polymerization pushing or pulling forces at the cortex or in the cytoplasm appear to be dispensable for repositioning spindles to the cell center.

**Figure 2.**
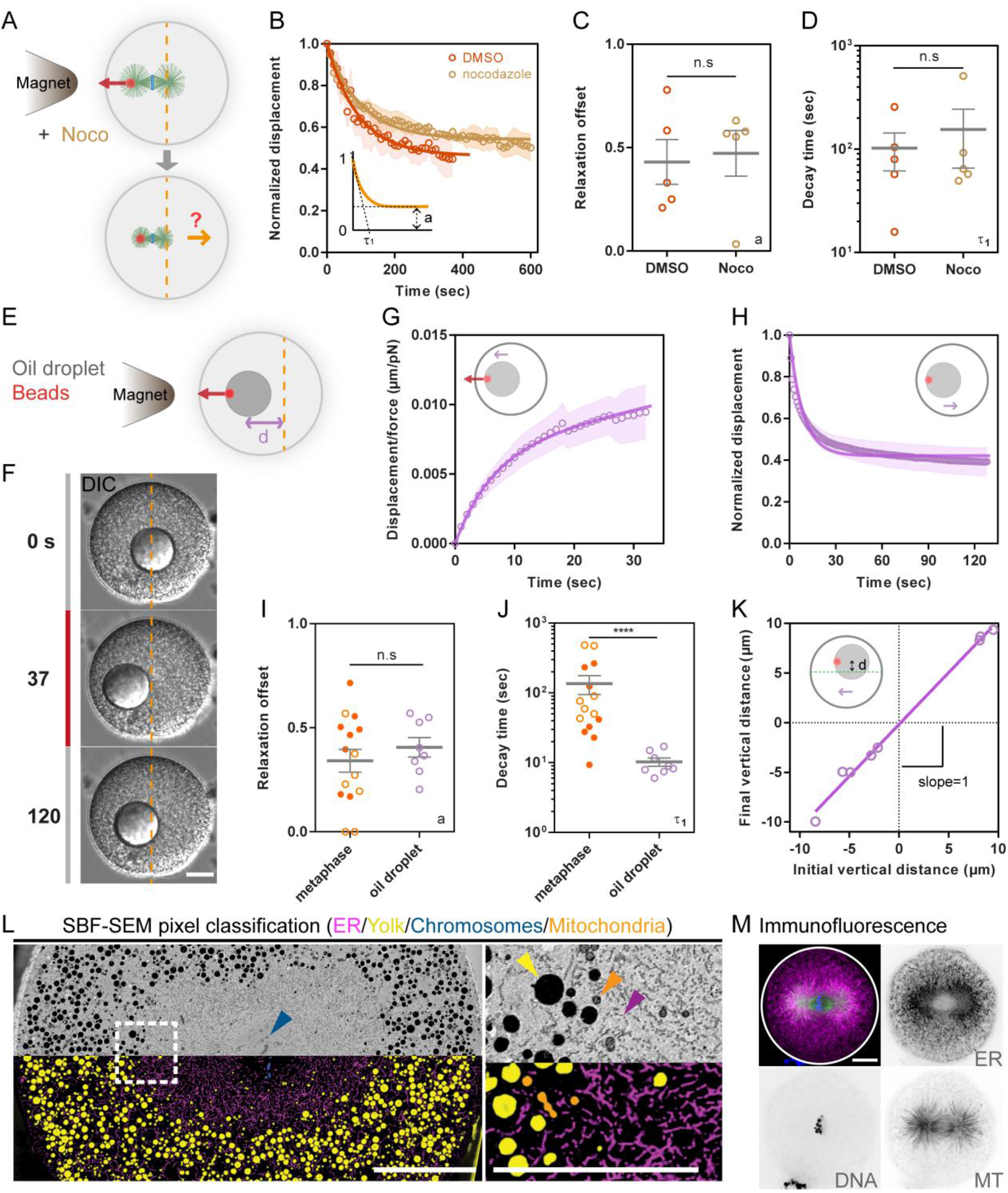
Visco-elastic restoring forces on spindles are associated to the material properties of bulk cytoplasm. **(A-B)** Metaphase spindles are displaced with magnetic tweezers and treated with either DMSO as controls (n=5) or Nocodazole to affect microtubules (n=5 cells), and their recoiling behavior to the cell center is assayed. **(C-D)** Quantification of the positional offsets to the cell center, a, and decay time-scales τ_1_ of relaxation curves plotted in B, using a single exponential model as indicated in the inset in B. **(E)** Scheme representing injected oil droplets containing hydrophobic magnetic beads used to actuate large objects with magnetic tweezers in the cytoplasm. **(F)** Time-lapse of a magnetic oil droplet displaced with an external magnetic force and recoiling upon force cessation. **(G)** Time evolution of the displacement measured from the initial position of the droplet and normalized by the applied force (n=5 cells). **(H)** Time evolution of the normalized displacement of the droplet back to its initial position upon force cessation. **(I-J)** Relaxation offset and decay time-scales for metaphase spindles (n=15) and oil droplets (n=8). **(K)** Final vertical offset plotted as a function of the initial vertical offset for droplets displaced horizontally with magnetic tweezers and let to relax (n=8). The line is a linear fit with a slope of 1.04. **(L)** Mid-section of a fixed metaphase zygote imaged with serial block face scanning electron microscopy (SBF-SEM) and corresponding pixel classification of different endomembrane compartments in the cytoplasm, and close up view at the border between the spindle and the rest of the cytoplasm. **(M)** Projected spinning disk confocal image of a metaphase zygote fixed and stained for Microtubules, DNA and Endoplasmic reticulum. Error bars and shades represent +/-S.E.M and +/-S.D/2 respectively. Results were compared using a two-tailed Mann–Whitney test. n.s, P > 0.05, ****, P < 0.0001. Scale bars, 20 µm and 10 µm in the close up view of 2L.

To more directly establish that viscoelastic repositioning forces are independent of spindle associated cytoskeletal elements, we sought to recapitulate early micro-rheology assays performed by Crick or Hiramoto (6-8), but using objects that have similar sizes as mitotic spindles. Inspired by recent experiments performed in mouse oocytes and in Xenopus extracts (27, 38), we embedded hydrophobic magnetic beads in soybean oil, and injected large 30-35 µm oil droplets to move them in the cytoplasm with magnetic tweezers. We purposely used unfertilized eggs, to circumvent the presence of large asters or spindles that could affect droplet motion in the cytoplasm from steric hindrance or by generating active flows and stresses

(39) (Fig 2E-2F). In the absence of external forces, oil droplets were immobile in the cytoplasm for long durations of up to 1h, much like female nuclei in these unfertilized eggs (Fig S2F-S2K). Remarkably, droplets exhibited a viscoelastic response to external forces similar to spindles, with a rapid initial constant velocity followed by a saturating elastic regime. Upon force cessation, droplets moved back towards their initial position with similar offsets as spindles, but shorter time-scales. Droplets viscoelastic recoils occurred along the same straight path as during force application, indicating that this elastic recoiling behavior is not restricted to objects initially positioned at the cell center (Fig 2G-2K and Movie S4). These data suggest that elements in bulk cytoplasm may generate viscoelastic reactive forces that move spindles or similar-sized passive objects back to their initial positions.

While mitotic spindles are often pictured as polar networks made of MT filaments, the accumulation of membranous organelles or other nuclear intermediate filaments on their MT network has suggested the existence of a spindle matrix, which could render them more physically akin to an impermeable droplet (15, 40). Accordingly, by performing Serial Block Face electron microscopy, we found that spindles were covered by packed endomembranes, with an “onion peel arrangement” typical of mitotic endoplasmic reticulum (ER) networks (41) (Fig 2L and Movie S5). This endomembrane accumulation is readily evident in DIC images as a smooth area around spindles (Movie S1-S3). Immunofluorescence further validated this accumulation, and segmentation of ER membranes provided an estimate of an upper-bound pore size of 0.2-0.5µm between membranes (Fig 2L-2M). Thus, metaphase spindles may be impermeable to relatively large objects and networks, a property which like oil droplets, allows them to be dragged by viscoelastic flows and forces from bulk cytoplasm.

### The cytoplasm applies large elastic and viscous drags to the mitotic spindle

To quantify restoring stiffness and viscous drags, we fitted experimental data with a three-element Jeffreys’ model, which provided the simplest 1D linear model for the observed rising and relaxation curves (42). This model has been employed to describe, among others, the rheology of suspensions of elastic spheres, polymeric liquids, and was previously used to understand viscoelastic properties of the cytoplasm (7, 43, 44). This model features an elastic spring of stiffness κ, in parallel with a dashpot of viscosity γ_1_, in series with a second dashpot of viscosity γ_2_, defining two characteristic time-scales. The first one, τ_1_ = γ_1_/k is the time-scale needed for spindle associated flows to charge the elastic elements in the material. The second, τ_2_ = γ_2_/k sets the rate of plastic yield of these elements or fluidization of the material, which limit elastic restoration and generates an offset in the relaxation (42) (Fig 3A-3B).

**Figure 3.**
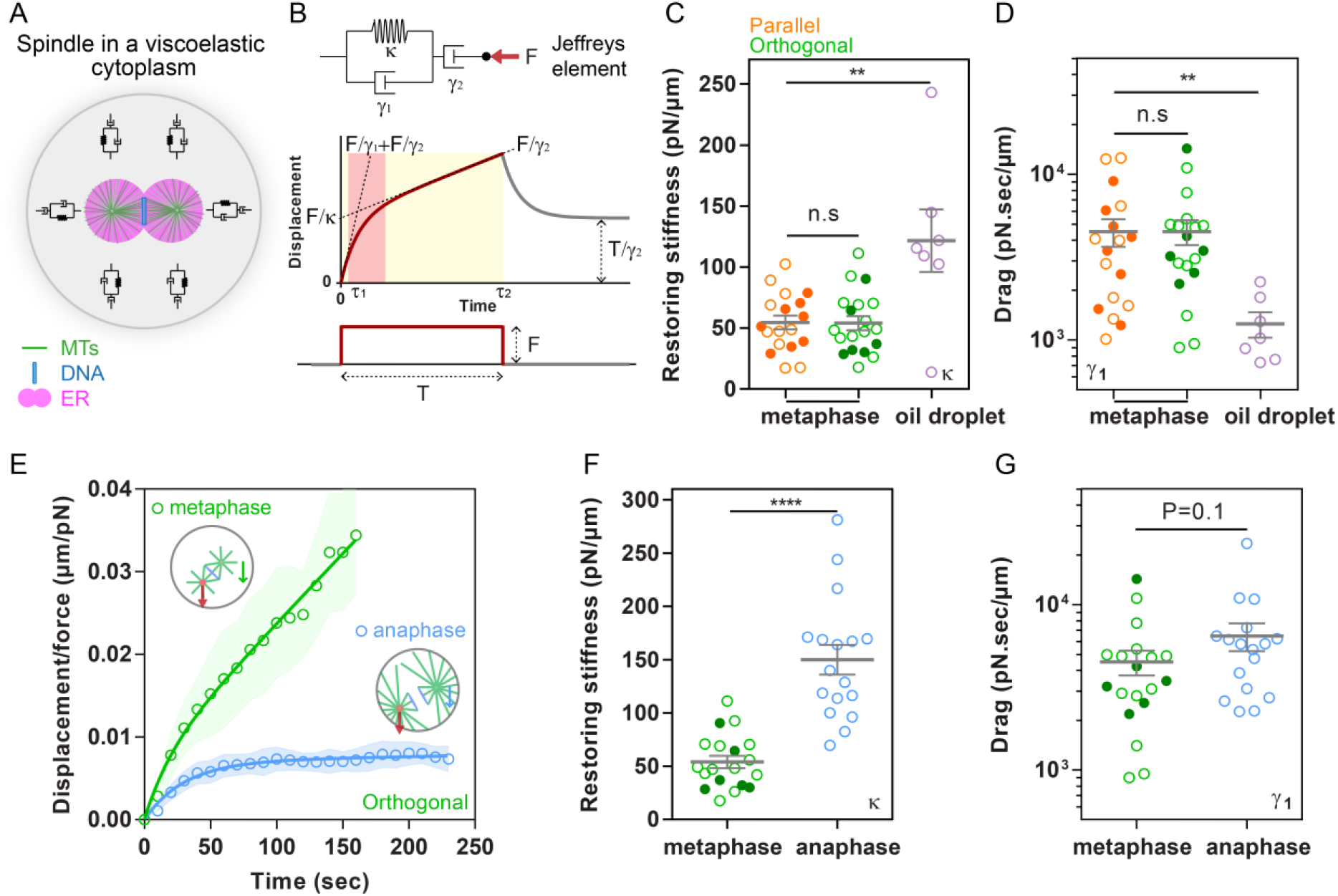
Cytoplasm material generates large elastic and viscous drags on mitotic spindles of similar magnitude as those applied by active microtubule arrays. **(A)** Scheme of metaphase spindles held in the cell center by viscoelastic elements in the cytoplasm. **(B)** Scheme of Jeffreys 3-element model made of a spring and a dashpot in parallel and another dashpot in series, and typical creep and relaxation behavior of this element. **(C-D)** Cytoplasm restoring stiffness and viscous drags on mitotic spindles moved parallel or orthogonal to the spindle axis (n= 18 and 19 respectively) and for oil droplets (n=7), computed using fits to the Jeffreys’ models represented in B. Open and closed dots correspond respectively to data obtained in control or MG132 treated cells. **(E)** Time evolution of the displacement measured from the initial centered position of the spindle and normalized by the applied force in metaphase and anaphase. **(F-G)** Restoring stiffness and viscous drags of metaphase and anaphase spindles moved orthogonal to the spindle axis (n=19 and 17 respectively). Error bars and shades correspond to +/-S.E.M and +/-S.D/2 respectively. Results were compared by using a two-tailed Mann–Whitney test. n.s, P > 0.05, **, P < 0.01, ****, P < 0.0001.

Using this model, we computed a restoring stiffness for spindles pulled along their long axis of κ = 55 +/-24 pN/µm (n=19 cells) and 54 +/-25 pN/µm for orthogonal pulls (n= 21 cells). This shows that a displacement away from the cell center of 5% of the cell diameter (∼4 µm), generates a restoring force of ∼250 pN. These are forces equivalent to that generated by hundreds of molecular motors (45). The stiffness measured for oil droplets was twice higher than for spindles, accounting for their shorter relaxation time-scales (Fig 3C and Fig 2J). One possible interpretation is that oil droplets are actuated in an unfertilized cytoplasm previously reported to be twice stiffer than the mitotic cytoplasm (7, 8). An alternative, is that the spindle may be porous to smaller elastic elements in the cytoplasm. These numbers amount to a lower-bound value of the effective bulk modulus, G=κ/6πR, with R the size of spindles, to be on the order of 0.2-0.3 Pa. This is smaller, but in the same range as previous rheological measurements of cytoplasm stiffness in different cell types and extracts (8, 10, 46).

Spindle viscous drags were unexpectedly high with a parallel drag of 4493 +/-3592 pN.s/µm and an orthogonal drag of 4498 +/-3359 pN.s/µm (Fig 3D). Considering reported values of cytoplasm viscosity in these cells (7), they amount to the drag of an object typically ten times larger than the spindle. This enhanced drag could be explained by the hydrodynamic coupling of the spindle with the cell surface which reduces spindle mobility (47, 48). Accordingly, theoretical predictions for a solid object moving in a viscous fluid contained in a sphere, with a radius twice that of the object, yield to a 10-20 fold increase of the object’s drag coefficients (49). This suggests that spindle large drags could primarily result from its interaction with the cytoplasm fluid confined by cell boundaries. Overall, these analyses support that the response of spindles to external forces are primarily associated to the viscoelastic properties of the cytoplasm (Fig S3A-S3D).

To compare the efficiency of bulk cytoplasm viscoelastic properties to that of active MT asters and motors, we next measured centering stiffness in anaphase, which follows metaphase by only few minutes. In anaphase, asters regrow to fill the whole cell volume, with an estimate of several hundreds of MTs contacting the cortex (Fig 1C and Fig S1E). Anaphase is thought to imply the largest MT based forces along the cell cycle, as MTs engage with dynein motors at the surface or in the cytoplasm, to separate chromosomes and move asters apart (26, 39, 50). Here, magnetic forces were only applied orthogonal to the spindle axis, as parallel pulls interfered with chromosome force-separation systems (Fig S3E-S3G). While the drags of anaphase spindles were similar to that found for metaphase spindles, a net difference was obtained in term of elastic behavior, with a restoring stiffness for anaphase spindles of 150 +/-57 pN/µm, nearly three times higher than during metaphase (Fig 3E-3G). These results suggest that bulk cytoplasm restoring forces can amount to ∼30% of the maximum MT/dynein based force-generating system in these cells.

### Cytoplasm forces depend on crowding and bulk F-actin meshworks

Crowding agents in bulk cytoplasm, that contribute to set elastic and viscous properties may include among others, cytoskeletal networks, endomembranes like Yolk granules and mitochondria, or ribosomes which are highly abundant in the cytoplasm (Fig 2L). As a general assay to affect cytoplasm crowding, we immersed cells ∼5 min prior to mitosis in diluted or concentrated artificial sea water (ASW) and pulled spindles at metaphase onset. We limited the amplitude of these shocks to a range in which spindles length, anaphase and cytokinesis were unaffected, but we noted delays in metaphase in hypertonic treatments (Fig S4A-S4D). A hypotonic treatment in 80% ASW caused water to flow into the cells with a minor cell size increase, and reduced spindle restoring stiffness and drags in this diluted cytoplasm to 50.2% and 49.6% of control values respectively. Conversely, a hypertonic treatment in 110% ASW, shrank cells by ∼6% and concentrated the cytoplasm, increasing spindle restoring stiffness and drags to 151.4% and 114.4% of control values, respectively (Fig 4A-4D and Fig S4E-S4H). These results demonstrate that viscoelastic reactive forces applied on spindles are directly related to cytoplasm crowding.

**Figure 4.**
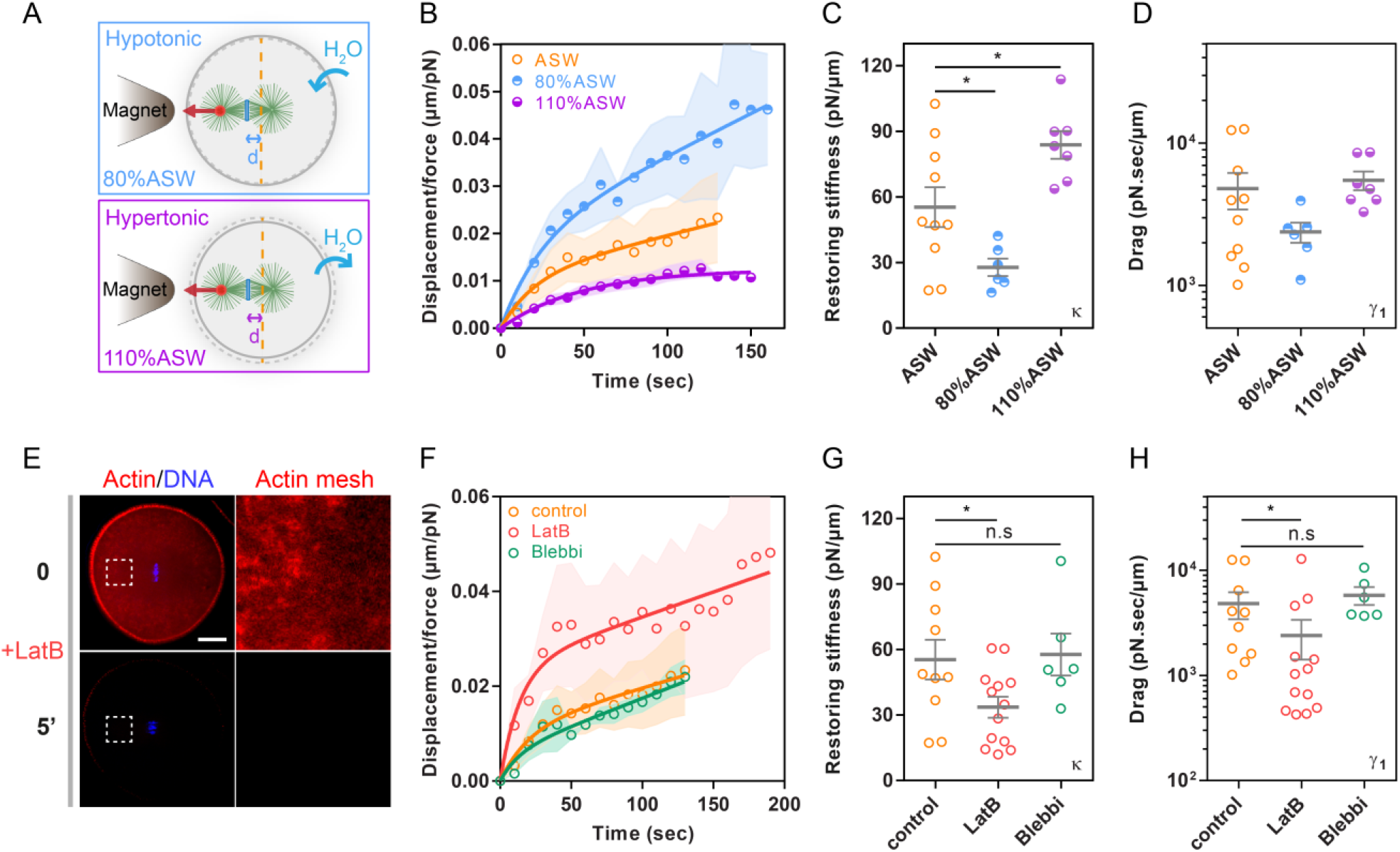
Influence of cytoplasm crowding and F-actin meshwork on spindle restoring stiffness and drags. **(A)** Scheme representing experiments to change cytoplasmic density by placing eggs in hypotonic or hypertonic artificial sea water (ASW). **(B)** Displacement normalized by applied force of metaphase spindles in control (n=10), hypotonic (n=6) and hypertonic (n=7) sea water. **(C-D)** Cytoplasm resorting stiffness and viscous drags on mitotic spindles moved parallel to the spindle axis, in controls, hypotonic or hypertonic sea water. **(E)** Confocal images of eggs injected with Utrophin-488 to visualize F-actin at metaphase in bulk cytoplasm (top) and ∼5 min after treating the same cell with Latrunculin B (bottom). The two images are treated with the same contrast to compare F-actin signals. **(F)** Displacement normalized by applied force of metaphase spindles in control (n=10), cells treated with Latrunculin B (n=13) and cells treated with Blebbistatin (n=6). **(G-H)** Cytoplasm restoring stiffness and viscous drags in controls, cells treated with Latrunculin B and cells treated with Blebbistatin. Error bars and shades correspond to +/-S.E.M and +/-S.D/2 respectively. Results were compared by using a two-tailed Mann–Whitney test. n.s, P > 0.05, *, P < 0.05. Scale bars, 20 µm.

Furthermore, as one element which has been shown to contribute to bulk cytoplasm material properties we tested the role of F-actin (10, 46). Imaging injected labelled utrophin, that binds F-actin, we detected a significant amount of disordered bulk F-actin meshes that surrounded the spindle, though unsurprisingly, the cortex was the most abundant part of the cell. Similar results were obtained with phalloidin staining (Fig 4E and Fig S4I). To affect F-actin, we treated cells with Latrunculin B, which disassembled F-actin within minutes, causing the cell surface to soften and shrivel (Fig 4E and Fig S4J-S4K). In Latrunculin B treated cells, spindle repositioning stiffness was 34 +/-17 pN/µm, 1.6 times lower than controls, and viscous drags were 2 times lower, more reduced than elastic values. As a consequence, the time-scales for both rising and relaxing phases in Latrunculin B were shorter than in controls, so that the re-centering dynamics, converged onto a similar positional offset, but occurred faster (Fig 4F-4H and Fig S4L-S4O). Importantly, this effect did not implicate putative acto-myosin contractile forces, as treatment with Blebbistatin to inhibit Myosin II did neither alter restoring stiffness and drags nor spindle repositioning dynamics (Fig 4F-4H and Fig S4I, S4L-S4O). These data suggest that bulk F-actin meshworks may act as important crowders that contribute to ∼40-50% of cytoplasm elastic and viscous drags on pulled spindle.

### Functional implications of cytoplasm mechanics for asymmetric and oriented divisions

We next investigated an important feature of spindle response, which is the linear rising phase at long time-scale and the offset in the relaxation, which reflect progressive plastic rearrangements or fluidization of elastic elements in the cytoplasm. A prediction of this effect is that a longer force application should dissipate more elastic energy, allowing spindles to stay further from their initial position when the force is released. Accordingly, in the Jeffrey’s model, the relaxation offset, *a*, depends mostly on the ratio of τ_2_ to the duration of force application, T, with 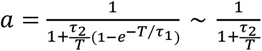 for sufficiently large T (Fig 5A). To test this, we arrested cells in metaphase with MG132, pulled spindles and held them for increasing periods of time (Fig 5B). Practically, this implied decreasing the pulling force gradually by progressively distancing the magnet, to avoid spindles moving onto the cortex. Remarkably, spindles held longer progressively lost their recoiling behavior with a well-matched alignment of experimental data on the predicted theoretical curve (Fig 5C-5D and Fig S5A-S5D). Therefore, elastic restoration by the cytoplasm may be very effective at short-time scales to stabilize spindles against random thermal or active forces, but vanishes over longer time-scales to facilitate spindle decentration during asymmetric divisions.

**Figure 5.**
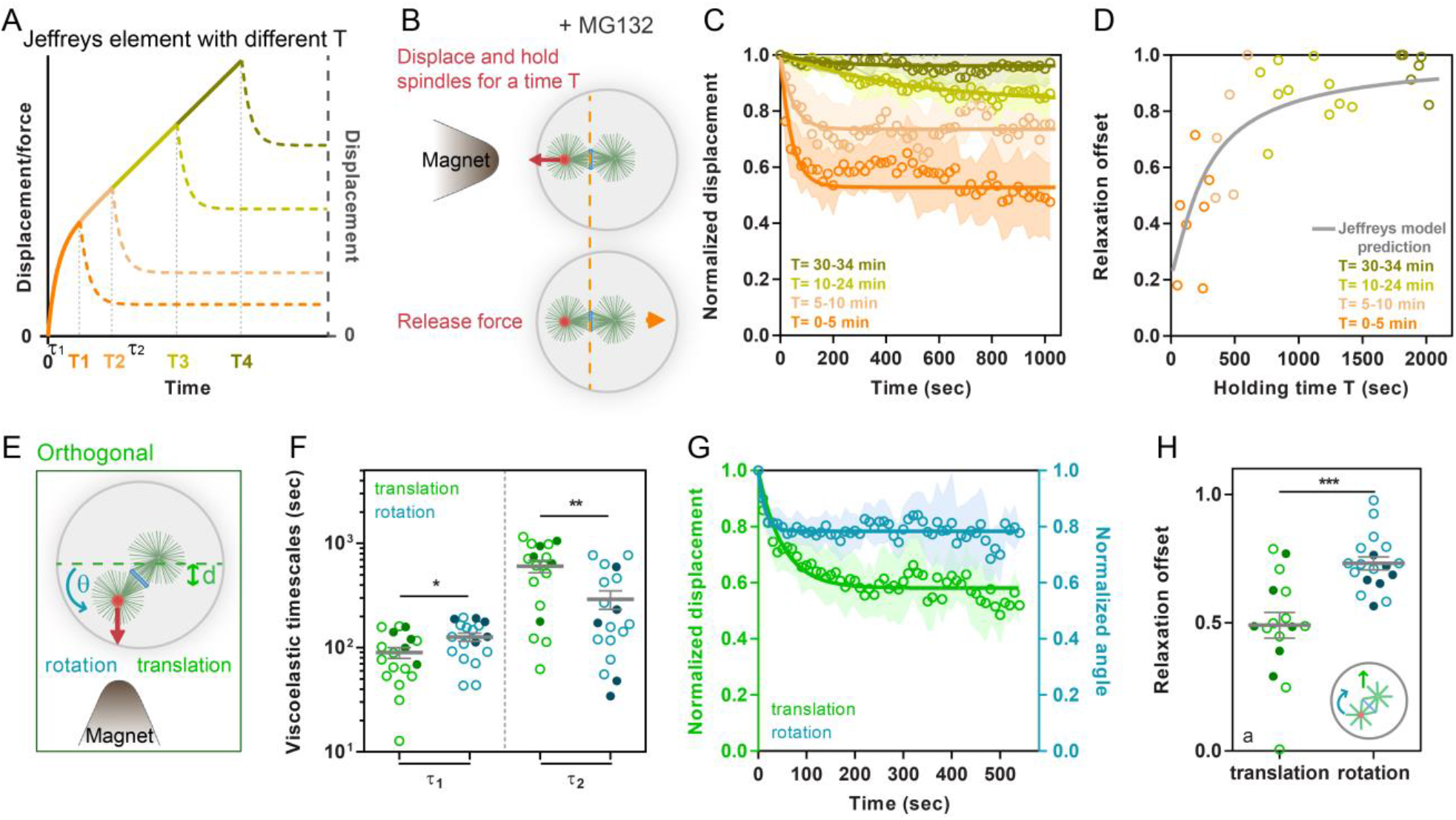
Dissipation of stored elastic energy and implications for asymmetric and oriented divisions. **(A)** Theoretical curve of creep and relaxation curves for different force application times in the Jeffreys’ model. Note how longer force applications tend to dissipate positional memory during relaxation. **(B)** Scheme explaining the assay to test the influence of force application time on relaxation. **(C)** Relaxation responses of metaphase spindles pulled and held with magnetic tweezers for increasing durations. **(D)** Relaxation offset, *a*, plotted as a function of force application duration, overlaid with the predicted theoretical curve form Jeffreys’ model, 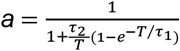, using measured values of τ_1_ and τ_2_. **(E)** Scheme representing how spindle rotation and displacement can be compared in the orthogonal pulling assay. **(F)** Viscoelastic time-scales τ_1_ and τ_2_ for rotational and translational spindle motions under force for the same cells and spindles (n=18). Open and closed dots correspond respectively to data obtained in control or MG132 treated cells. **(G)** Rotational and translational relaxation curves upon force cessation. **(H)** Relaxation offset for rotational and translational relaxation. Error bars and shades correspond to +/-S.E.M and +/-S.D/2 respectively. Results were compared by using a two-tailed Wilcoxon test. *, P < 0.05, **, P < 0.01, ***, P < 0.001.

As another important read out of this effect, we found that τ_2_ was significantly smaller in rotation than in translation (Fig 5E-5F and Fig S5E-S5H). This was reflected both in the more linear shape of the rising curve under force and in the higher values in the angular relaxation offset as compared to translation for the same spindles (Fig 1L and Fig 5G-5H). Thus, rotating spindles appear to dissipate faster rotationally stored elastic energy, causing spindle reorientation to be less-well restored by the cytoplasm than translation. This effect will tend to favor spindle rotation over translation, allowing for instance active force generators to easily tilt spindles, without translating it. We conclude that the dissipation of spindle elastic restoration by plastic rearrangements or fluidization of the cytoplasm and their time-dependence may be highly relevant to understand spindle positioning phenotypes.

### Large-scale flows of cytoplasm material associated with spindle motions

To understand how the cytoplasm reorganizes in response to spindle motion and forces, we mapped cytoplasmic flows. We tracked cytoplasmic granules of ∼1 µm in size with Particle Image Velocimetry (PIV) (movie S6). These elements are larger than measured spindle pore size and are dominated by advection with estimated Peclet numbers exceeding 10^2^-10^3^, therefore best representing potential rearrangement of entangled elastic meshworks in the cytoplasm. When spindles were pulled along their long axis, the cytoplasm flowed along with the spindle and recirculated, forming large symmetric vortices mirrored along the spindle axis. As spindles relaxed, similar vortices formed but with an opposite rotational direction (Fig 6A-6B and Fig S6A). By computing the local divergence of the flow pattern, which provides a qualitative indicator of how a viscoelastic material may contract or expand, we found that the portion of the cytoplasm at the front of a pulled spindle appeared compressed, while the portion at the back was more stretched (Fig S6D-S6E). Vorticities in the flow field allowed to visualize shear, and to extract a time-scale from the inverse of the shear rate, on the order of 570 +/-260 s, close to the time scales for plastic dissipation τ_2_ of spindle viscoelastic response (Fig S6B, S6G and S6K). These data indicate that spindle displacement causes the viscoelastic cytoplasm to locally compress and/or stretch and apply reactive elastic forces, and shear away from the spindle front.

**Figure 6.**
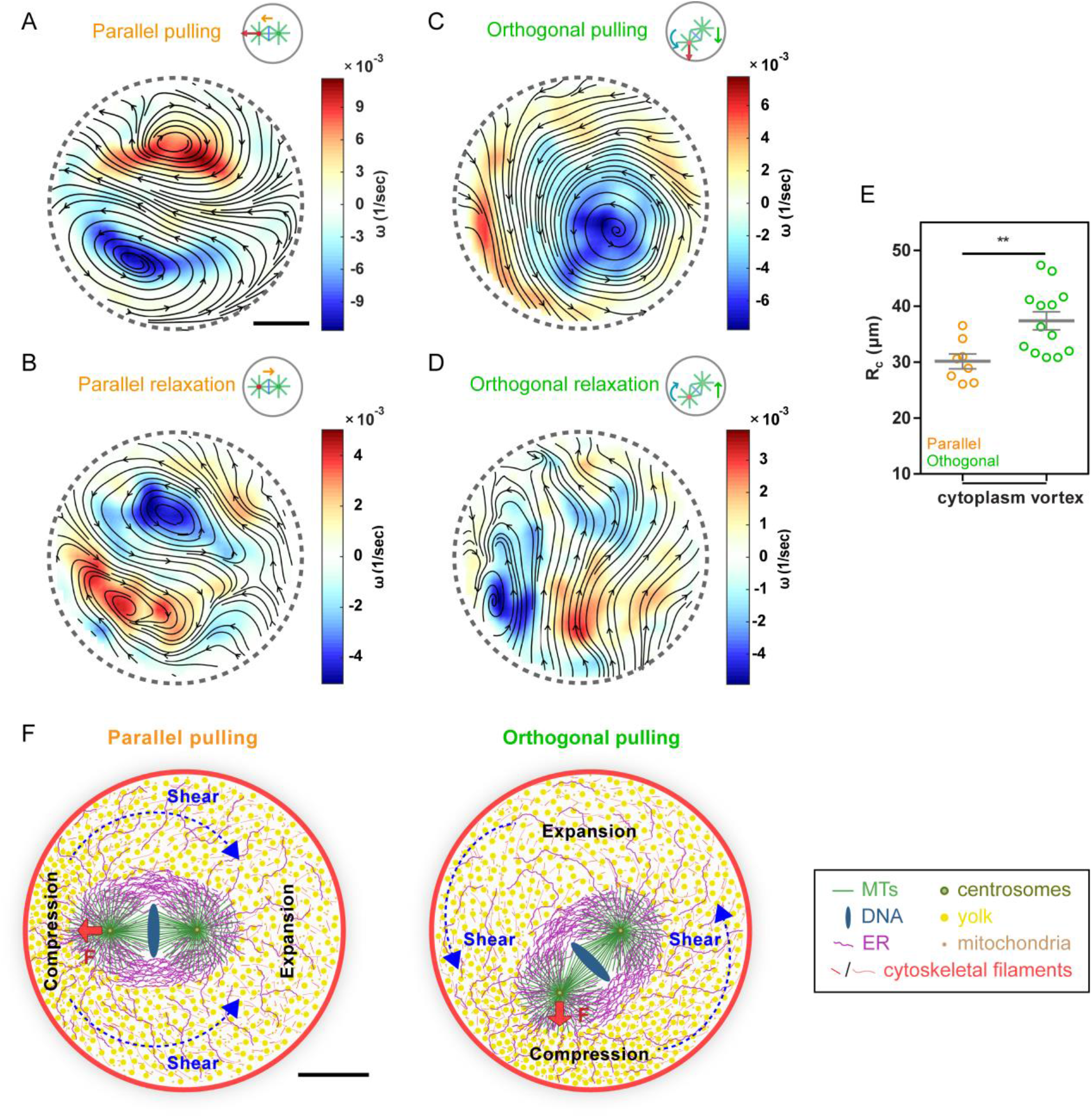
Large-scale flows of cytoplasm elements associated with spindle motions. **(A-D)** Cytoplasm streamlines averaged on a duration of 60 s and size of 50 µm during parallel force application on spindles (A) and during spindle recoils (B), or during orthogonal force applications averaged on a duration of 100 s and size of 90 µm (C) and relaxation (D). Streamlines are superimposed onto a color map of local flow vorticities, ω. **(E)** Quantification of flow vortices size during parallel or orthogonal pulls (n=8 and 13 cells respectively). **(F)** Proposed models based on flow map analysis and force response of metaphase spindles. In parallel pulls, the cytoplasm made of relatively large objects including cytoskeletal elements and endomembranes (Yolk, mitochondria or ER), is compressed at the spindle front and stretched at the rear, which causes it to push or pull back the spindle with elastic restoring forces. At the same time, the cytoplasm material is sheared along the spindle. In orthogonal pulls, compression of the cytoplasm may be prominent at the spindle front, like in parallel pulls, but in rotation, the broken symmetry causes shear to dominate over compression. Shear stresses fluidize or plastically rearrange the cytoplasm faster than in compression potentially accounting for reduced elastic restoring of spindle angles as compared to positions upon force applications. Results were compared by using a two-tailed Mann–Whitney test. **, P < 0.01. Scale bars, 20 µm.

In orthogonal pulls, the flow patterns were markedly different, with one dominant large vortex centered around a point located bottom-right in the cell with respect to the pulled spindle pole (Fig 6C and Fig S6F). This vortex was significantly larger than in translation (Fig 6E and Fig S6C, S6H). Furthermore, the divergence map appeared asymmetric orthogonal to the spindle axis, but did not reveal notable left-right asymmetric patterns, plausibly reflecting the reduced ability of the cytoplasm to rotate back spindles (Fig S6I-S6J). In relaxation, the flows were mostly vertical, in agreement with the dominance of a translational over rotational spindle recoil (Fig 6D). Therefore, a spindle rotating in the confined boundaries of a cell, appears to shear rather than compress the cytoplasm (Fig 6F). In light of the differential viscoelastic response of translating vs rotating spindles, these flow analyses suggest that shear fluid stress could cause cytoplasm elements to plastically yield faster than in compression, a process typical of many viscoelastic materials (42). We conclude that the organization of viscoelastic flows in response to spindles moving in a cytoplasm confined by cell boundaries, may have a key impact on mitotic spindle positioning phenotypes (Fig 6F).

## Discussion

### Function of bulk cytoplasm rheology in the mechanics of spindle positioning

How spindles are positioned and oriented in embryos and tissues is a fundamental problem for cell and developmental biology highly relevant to the emergence of developmental disorders and cancer (14, 51). One current dogma in many animal cells, is that spindles are placed with respect to cell boundaries by active forces from astral MTs that grow to the cell cortex. Here, by exploring a regime where mitotic MT asters do not reach out to the cortex, we demonstrate that the cytoplasm acts as a viscoelastic medium that holds spindles or other large objects in place, and moves them back if their position is perturbed. Restoring forces are large and shall participate in the force balance positioning spindles and asters, by for instance opposing asymmetric cytoskeleton forces during nuclear or spindle decentration for asymmetric divisions, or those needed to center asters at fertilization (20, 37, 48). Accordingly, the cytoplasm restoring stiffness on spindles measured here, is of similar magnitude as that measured during sperm aster centration in the same model system (37), and ∼3X higher than that associated to metaphase spindle maintenance at the cell center in *C. elegans* (26). Our findings also suggest that restoring forces vanish faster in rotation than in translation, which we attribute in part to the geometry of rotational shear flows, and anisotropies in plastic yields of the cytoplasm material. Therefore a tilt acquired during force exertion is better maintained than a positional offset. This could allow an asymmetric cortical domain enriched in motors to reorient spindles without creating an asymmetric division, a phenotype commonly observed in many tissues (14, 52, 53).

It is important, however, to outline that at rest, the cytoplasm does not apply any net force, and cannot *a priori* center or decenter spindles, asters or nuclei. In large cells, this is achieved earlier in the cell cycle by interphase MT asters that reach to the cell boundaries, and pre-position centrosomes before metaphase (30, 31). We propose that the cytoplasm holds these prepositioned centrosomes in place. As spindles scale with cell size, cytoplasm rheology could very well be also relevant to spindle positioning in smaller cells lacking proper mitotic asters, or in which astral MTs do not reach the cell surface (54, 55). However, an important open question, is how much these properties contribute in cells where numerous MTs clearly contact the cortex to exert forces that stabilize asters or spindles (26, 37). One possibility is that the impact of cytoplasm elasticity is reduced as astral MTs and motors push back organelles and networks away from spindles, effectively confining elastic elements to a zone much closer to the cortex. Another one, which we favor, is that transport by motors along MTs could fluidize the cytoplasm space occupied by mitotic asters, effectively decreasing the elastic response of the cytoplasm. Further work will be required to understand how active polar cytoskeleton forces and material properties of bulk cytoplasm are integrated to regulate cell division and organization.

### Probing cytoplasm material properties at the scale of mitotic spindles

The cytoplasm is a complex material, whose rheological properties and their impact on cellular functions are emerging as important concepts in cell biology. Here, we probed cytoplasm rheology at scales and speeds, typical of physiological mitotic spindle repositioning and reorientation. In line with previous rheological descriptions of the cytoplasm for small objects (7, 8), we find that a simple linear 1D viscoelastic Jeffreys’ model can predict key rheological behavior including rising and relaxation curves, and time-dependent elastic dissipation. Although this suggests that we can treat the effect of the cytoplasm as that of an “inert” Jeffreys’ material, these physical properties, are likely dictated and regulated by energy-driven active metabolic processes and random motor motion (10, 12). We also envision that more advanced non-linear viscoelastic or poroelastic description of the cytoplasm could capture many of the features we report in here. Elasticity could for instance be graded, being stiffer close to the cortex and softer in the cell interior, due to the progressive stacking of organelles and networks. Inspections of electron micrographs do not immediately support this view, at least in this system. Poroelasticity would picture the cytoplasm as a porous elastic solid, with a small pore size bathed in cytosolic liquid (3, 56). As spindles would compress this poroelastic medium, slowly dissipating pressure gradients would form and push back spindles in place.

Beyond physical frameworks to describe cytoplasm rheology, another important question is which elements define relevant material properties of the cytoplasm at the spindle scale. Meshworks of bulk F-actin filaments contributed to a fraction of ∼40-50% of viscous and elastic behavior, acting as one important crowding agent and organelle stabilizer at this scale (57, 58). Intermediate filaments, like keratin or vimentin could not be tested here, but have been proposed to influence elastic properties of egg cytoplasm extracts (46). In addition, we propose that small organelles like mitochondria, yolk granules or lysosomes could behave as dense suspensions of small elastic colloids, which generate mesoscopic elasticity as is known for classical suspension such as paints or bitumen (43). Further experiments of cytoplasm reconstitution will be required to determine which of these elements are dominant or dispensable at the spindle scale, or assay the role of their potential interactions.

Finally, our experiments comparing spindles and oil droplets, raise interesting questions on the physical nature of the mitotic spindle itself. In the context of our assays, we propose that spindles behave as objects impermeable to components larger than a fraction of a micron. Recent reports suggest that small particles of ∼10nm in size can diffuse in and out of spindles (59). Spindles may thus be largely porous to proteins and molecular complexes such as ribosomes, so that elastic and viscous forces they experience may only result from the response of larger organelles and networks in the cytoplasm. Mitotic spindles also feature important elastic properties, with axial stiffness measured *in vitro* to range around 80-900 pN/µm (60). In our experiments, we detected minor lengthening of mitotic spindles under force, which amounted to similar values. Recent experiments on MT asters also support models of more porous or elastic structures with important implications for mechanisms of centrosome movement (39). By bringing numbers which have been largely missing in the literature, force measurements within live cells will strongly impact our understanding of the very basic mechanisms of cell organization and morphogenesis.

## Acknowledgment

We thank P. Lenart, B. Lacroix and J. Dumont for sharing reagents, as well as our colleagues C. Leduc, S. Dmitrieff, G. Romet-Lemonne, Y. Bellaïche and J-L. Maître for carefully reading this MS. We gratefully thank all members of the Minc team for discussion and technical help. J.X. acknowledges the “École Doctorale FIRE - Program Bettencourt”, a fellowship from the Chinese Scholarship Council (201708070046) and from the LabEx “Who am I?” (ANR-11-LABX-0071). J.N. is supported by a fellowship from the ARC foundation (PDF20191209818) and is an EMBO Non-Stipendiary Fellow (ALTF 881-2019). We acknowledge the ImagoSeine core facility of the Institut Jacques Monod, member of Infrastructure en Biologie Santé et Agronomie and France-BioImaging (ANR-10-INBS-04) infrastructures. This work was supported by the Centre National de la Recherche Scientifique (CNRS), the Université de Paris, and grants from La Ligue Contre le Cancer (EL2021.LNCC/ NiM), the Agence Nationale pour la Recherche (ANR, “TiMecaDev”), the Fondation Bettencourt Schueller (“Coup d’élan”), and the European Research Council (ERC CoG “Forcaster” no. 647073) to N.M.

## Material and methods

Detailed methods are provided in the SI appendix. They include protocols to obtain and handle sea urchin animals and gametes, and procedures used for chemical inhibitions of cytoskeletal components and modulations of sea water osmolarity. All methods for live and fixed imaging of zygotes are also provided, including those used for transmission electron microscopy, serial block face electron microscopy and endomembrane registration in three dimensions. Finally, we also provide all detailed procedures for injecting magnetic beads, calibrating magnetic forces *in vitro*, and computing magnetic forces *in vivo*, as well as extract viscoelastic parameters and map and analyze viscoelastic flows in live cells.

