## Supplementary Information for "Contribution of cytoplasm viscoelastic properties to mitotic spindle positioning"

### SUPPORTING INFORMATION

This supporting information file contains Supplemental methods, 6 Supplemental Figures and Figure Legends and supporting movie legends.

### Supplemental methods

#### Sea urchin gametes

Purple sea urchins (*Paracentrotus lividus*) were obtained from Roscoff Marine station (France) and maintained at 16°C in aquariums of artificial sea water (ASW; Reef Crystals, Instant Ocean). Gametes were collected by intracoelomic injection of 0.5 M KCl. Sperm was collected dry and kept at 4°C and used within 1 week. Eggs were rinsed twice with ASW, kept at 16 °C, and used on the day of collection. Unfertilized eggs were transferred and adhered on protamine-coated glass-bottom dishes (MatTek Corporation) after removing the jelly coat through an 80-µm Nitex mesh (Genesee Scientific), injected and fertilized under the microscope.

#### Chemical inhibitors

Inhibitors were prepared in 100X stock aliquots in dimethyl sulfoxide (DMSO). To block metaphase, eggs were incubated for 30 min in 50 µM MG132 (Sigma-Aldrich) before fertilization. Nocodazole (Sigma-Aldrich) was applied ~3 min before spindle relaxation at a final concentration of 20 µM. Latrunculin B (Sigma-Aldrich) was added ~5 min before pulling the spindle at a final concentration of 20 µM. Blebbistatin (Sigma-Aldrich) was added 15 min post fertilization at a final concentration of 100 µM.

#### Hypotonic and Hypertonic shocks

To manipulate cytoplasm density with hypotonic or hypertonic shocks, ASW was prepared to 80% or 110% of its normal content, respectively, by adjusting the amount of DI water to salts mixtures. Eggs were rinsed with these different water at prometaphase, typically 5 min before force application. At these sea water concentrations, cells formed normal spindles and underwent a normal cytokinesis, but metaphase was longer in concentrated ASW. Much lower concentration of 50% ASW were also assayed, but led to large leaks of cytoplasm from the site of injection that dragged spindles out of cells. Higher concentration of 120% or 150% ASW led to cytokinesis failures.

#### Immunostaining

Immunostaining was performed using procedures described previously (1). Samples were fixed for 70 min in 100 mM Hepes, pH 6.9, 50 mM EGTA, 10 mM MgSO4, 2% formaldehyde, 0.2% glutaraldehyde, 0.2% Triton X-100, and 400 mM glucose. To reduce autofluorescence, eggs were then rinsed 3 times in PBS and placed in 0.1% NaBH4 in PBS freshly prepared, for 30 min. Eggs were rinsed with PBS and PBT (PBS + 0.1% TritonX) and blocked in PBT supplemented with 5% goat serum and 0.1% bovine serum albumin (BSA) for 30 min. Samples were rinsed with PBT before adding primary antibodies. For MT staining, cells were incubated for 48 h at 4°C with a mouse anti-α-tubulin (DM1A; Sigma-Aldrich) primary antibody at 1:5,000 in PBT, rinsed 3 times in PBT and incubated for 4 h at room temperature with anti-mouse secondary antibody coupled to Dylight 488 (Thermo Fisher Scientific) at 1:1,000 in PBT for 4-5h. For ER staining, samples were treated with a rabbit anti-KDEL (Invitrogen) primary antibody at 1:2,000 in PBT, for 48h, rinsed in PBT and incubated with an anti-rabbit secondary antibody coupled to Dylight 650 (Thermo Fisher Scientific) at 1:1,000 in PBT. To stain F-actin, samples were incubated for 1h in a solution of Rhodamine Phalloidin at 4U/ml in PBT. Eggs were washed three times in PBT then twice in PBS, transferred in 50% glycerol in PBS, and finally transferred into mounting medium (90% glycerol and 0.5% N-propyl gallate in PBS). Fixations of cells treated with Nocodazole and Blebbistatin were performed 10 min and 30min respectively after adding the drug (Fig S2B and Fig S4I).

**Fixation for electron microscopy**

To visualize astral MTs around spindles, one cell stage zygotes were fixed in metaphase (1h post-fertilization) in 0.4 M sodium acetate, 2% glutaraldehyde (Electron Microscopy Sciences) during 30 sec and in 0.4 M sodium acetate, 2% $OsO_4$ (Electron Microscopy Sciences) during 30min (Asnes and Schroeder, 1979), and transferred for 1h to 2% glutaraldehyde, 1% Osmium, then washed 15 min with the same buffer and 2 x 15 min in water. Dehydration was performed using graded concentrations of ethanol in water for 10 min each. Resin infiltration was performed by incubating 30 min in a 30% Agar low viscosity resin (Agar Scientific Ltd), then 30 min in a 50% Agar low viscosity resin, 30 min in a 75% resin followed by a 1h and overnight incubation in pure resin. The resin was then changed twice and the samples were incubated for 1 hour prior to inclusion in BEEM capsules and 18 hours polymerization at 60 °C. 70 nm sections were obtained using an EM UC6 ultramicrotome (Leica), and post-stained in 4% aqueous uranyl acetate followed by lead citrate.

For ER endomembrane visualization by Serial Block Face electron microscopy , one cell stage zygote were fixed in metaphase in 0.2 M cacodylate buffer, 0.25 M sucrose, 2% paraformaldehyde, 2% glutaraldehyde during 1h and progressively transferred in 0.2 M cacodylate buffer, 0.35 M NaCl, and stored at 4°C until processing. Samples were then prepared for Serial Block Face using the NCMIR protocol (https://ncmir.ucsd.edu/sbem-protocol). They were washed 3 times with cold cacodylate and post-fixed for 1 hour in a reduced osmium solution containing 1% osmium tetroxide, 1.5% potassium ferrocyanide in 0.15 mM sodium cacodylate buffer, followed by incubation with a 1% thiocarbohydrazide (TCH) solution in water for 20 min at room temperature (RT). Subsequently, samples were fixed with 2% $OsO_4$ in water for 30 min at room temperature, followed by 1% aqueous uranyl acetate at 4 °C overnight. The samples were then subjected to *en bloc* Walton's lead aspartate staining (*2*) at 60 °C for 30 min. Then, samples were dehydrated in graded concentrations of ethanol for 10 min each. The samples were infiltrated with 30% Agar low viscosity resin (Agar Scientific Ltd) for 1 hour, 50% Agar low viscosity resin for 2 hours and 100% Agar low viscosity resin overnight. The resin was then changed and the samples further incubated during 3 hours prior to inclusion in BEEM capsule Bottle Neck tips (EMS) and polymerized for 18h at 60 °C. The polymerized blocks were mounted onto special aluminum pins for SBF-SEM imaging (FEI Microtome 8mm SEM Stub, Agar Scientific), with two part conduction silver epoxy kit (EMS, 190215).

**Live-probe for the cytoskeleton**

TauMBD-mCherry constructs were expressed in BL21 (DE3)-RIL (Agilent) cells, purified on a Ni-NTA resin (Invitrogen) and dialyzed with PBS buffer using a 10000MWCO dialysis cassette (ThermoScientific). Tubulin-Atto565 was a gift from Benjamin Lacroix. Tubulin was purified from pig brains and labelled with NHS-ester-ATTO 565 (ATTO-TEC) (2). UtrCH-AlexaFluor488 was a gift from Peter Lenart. UtrCH recombinant protein was expressed in *E.coli*, purified on a Ni-NTA resin (Qiagen), and labelled with AlexaFluor 488 succinimidyl ester (Invitrogen) (3).

**Magnetic force application**

Magnetic tweezers were implemented as described previously (4, 5). The magnet probe used for force applications *in vivo* was built from three rod-shaped strong neodymium magnets (diameter 4 mm; height 10 mm; S-04-10-AN; Supermagnet) prolonged by a sharpened steel piece with a tip radius of ~50 µm to create a magnetic gradient. The surface of the steel tip was electro-coated with gold to prevent oxidation. The probe was controlled with a micromanipulator (Injectman 4, Eppendorf), mounted on an inverted epifluorescent microscope.

Super-paramagnetic particles (diameter 800nm; NanoLink; Solulink) with spontaneous minus end–directed motion were used to apply magnetic forces on the spindles (4, 5). To prepare beads for injection, a solution of 10 µl of undiluted streptavidin-beads was first washed in 100 µL of washing solution (1 M NaCl with 1% Tween-20), and sonicated for 5 min. The beads were then rinsed in 100 µl PBS, incubated 15 min in 100 µl 2 µg/ml Atto565-biotin (Sigma-Aldrich), rinsed again in 100 µl PBS, and finally re-suspended in 20 µl PBS and kept at 4°C until use. Unfertilized eggs were placed on a protamine-coated glass bottom dish. The bead solution was injected using a micro-injection system (FemtoJet 4, Eppendorf) and a micro-manipulator (Injectman 4, Eppendorf). Injection pipettes were prepared from siliconized (Sigmacote) borosilicate glass capillaries (1 mm diameter). Glass capillaries were pulled using a needle puller (P-1000, Sutter Instrument) and ground with a 40° angle on a diamond grinder (EG-40, Narishige) to obtain a 10 µm aperture. Injection pipettes were back-loaded with 2 µl of bead solution before each experiment, and were not re-used. After fertilization, beads were spontaneously transported along MTs and formed a large aggregate at the center of the aster. This aggregate stayed stably at the centrosome and occasionally split at centrosome duplication. Thus at metaphase, beads sometimes ended up on one or the two spindle poles (Fig S1H and S1I). The targeting of beads towards the centrosomes presumably occurred in a dynein- and microtubule-dependent manner. Thus in the presence of 20 µM Nocodazole, beads could detach from the spindle pole under external forces (Fig S2A, red arrowhead).

To image spindle displacement and rotation under force, unfertilized eggs stuck on protamine-coated glass dishes were incubated in 10 µg/ml Hoechst (Sigma-Aldrich) and subsequently injected with magnetic beads, and fertilized. At metaphase, chromosomes marked with Hoechst, lined up along the metaphase plate, and the spindle was easily visible as a dumbbell-shaped smooth region in DIC (Differential Interference Contrast), presumably because of its association with the ER (Movie S1). These DIC images allowed to select spindles that were planar and to define the long and short axis directions for force applications. The magnet was then rapidly moved close to the eggs and held at a fixed position either at a position along the spindle axis (for parallel pulls) or orthogonal to the axis for orthogonal pulls (see Movie S2-S3). The end of metaphase was captured as the first time-point when chromosome separated, and the end of anaphase as the first time-point of cleavage furrow ingression. For experiments implicating long force application (Fig 5), the magnet was progressively moved away from the egg using the automatic stage of the microscope.

To actuate oil droplets, unfertilized eggs were placed on protamine-coated glass-bottom dishes. To pull the oil droplets in the cytoplasm, a suspension of 10 µl of 1.2 µm hydrophobic superparamagnetic beads (magtivio, MagSi-proteomics C18) was washed in 100 µl of 30, 50, and 70 percent ethanol solution. It was then dried in vacuum for 20 minutes and re-suspended in 5 µl soybean oil (Naissance; Huile de soja). Preparation of glass capillaries and injection were done with the same methods as for other beads injection described above. All the injected eggs in each sample were surveyed to select the oil droplets with a sufficient amount of beads for pulling. After approaching the magnet, beads inside the oil formed aggregates and slowly moved toward the magnet while the oil droplet was stationary, until the aggregate contacted the oil-cytoplasm interface at the side facing the magnet tip. Large aggregates could not usually cross the interface due the oil surface tension, aggregate size, and hydrophobic properties of the beads, and were used to pull oil droplet in the cytoplasm. Upon force application, the magnet was quickly retracted when the oil distance to the cell cortex was ~10 µm, which caused the aggregate to detach from the oil/cytoplasm interface and the droplet to relaxed backward along the previous pulling force axis (Fig S2J-K).

**Imaging**

Time-lapses of spindles and oil droplets moving under magnetic force were recorded on two inverted microscope set-ups equipped with a micromanipulator for magnetic tweezers, at a stabilized room temperature (18–20°C). The first set-up was an inverted epifluorescence microscope (TI-Eclipse, Nikon) combined with a complementary metal–oxide–semiconductor (CMOS) camera (Hamamatsu), using a 20X dry objective (Apo, NA 0.75, Nikon) and a 1.5X magnifier, yielding a pixel size of 0.216 µm. The second one was a Leica DMI6000 B microscope equipped with an A-Plan 40x/0.65 PH2 objective yielding a pixel size of 0.229 µm, equipped with an ORCA-Flash4.0LT Hamamatsu camera. Both microscopes were operated with Micro-Manager (Open Imaging).

To visualize MTs or bulk F-actin with live probes, live imaging was performed on a spinning-disk confocal microscope (TI-Eclipse, Nikon) equipped with a Yokogawa CSU-X1FW spinning head, and an EM-CCD camera (Hamamatsu), using a 60X water-immersion objective (Apo, NA 1.2, Nikon), equipped with a 3D micromanipulator. Imaging of immunostained cells was performed on a confocal microscope (Zeiss, LSM980) coupled with an Airyscan 2 module in confocal mode with a 63X water immersion objective (NA, 1.4; C-Apo; Zeiss).

**Electron Microscopy**

Thin sections of eggs fixed as described above were observed by transmission electron microscopy (TEM) at 120 kV with a Tecnai12 transmission electron microscope (Thermo Fischer Scientific) equipped with a 4K Oneview camera (Gatan). For Serial Block Face scanning electron microscopy

(SBF-SEM), samples mounted on aluminum pins were trimmed and inserted into a TeneoVS SEM (Thermo Fisher Scientific). Acquisitions were performed with a beam energy of 2.7kV, a current of 400pA, in LowVac mode at 40Pa, a dwell time of 1µs per pixel and sections of 100nm. The pixel size was 20nm.

**Magnetic force calibration**

Magnetic forces were calibrated *in vitro* following procedures described previously (4, 5). The magnetic force field created by each magnet tip used was first characterized by pulling 2.8 µm mono-dispersed magnetic beads (Dynal) in a viscous test fluid (80% glycerol, viscosity 8.0×10−2 Pa sec at 22°C) along the principal axis of the magnet tip. Small motion of the fluid was subtracted by tracking 4 µm non-magnetic tracer fluorescent in the same solution. The speed of a magnetic bead V was computed as a function of the distance to the magnet, representing the decay function of the magnetic force, and fitted using a double exponential function.

To compute the dependence of the force on aggregate size, bead aggregates from the same beads as those used *in vivo* in sizes ranging from 2 to 8 µm, similar to that observed in cells, were pulled in the same fluid as above. The speed $V_a$ was measured and translated into a force using Stokes' law $F=6\pi\eta RV_a$, where η was the viscosity of the test fluid, R the aggregate effective radius defined using the longest length $L_1$ and the length perpendicular to the longest axis, $L_2$, as $R=\frac{1}{2}\sqrt{L_1L_2}$. The force–size relationship at a fixed distance from the magnet was well represented and fitted by a cubic function. These speed–distance and force–size relationships were combined to compute the magnetic forces applied to spindles inside cells as a function of time, from the size of aggregates at spindle poles and their distance to magnet tips.

Magnetic force calibration for oil droplets experiments followed the exact same procedure as above but establishing the force-size relationships for aggregates of hydrophobic beads used in oil droplets (Fig S2G). In live-cell experiments the sizes of the bead aggregates inside the oil droplets were measured at three different positions in the oil in the bright field, and were averaged.

**Analysis of spindle position, orientation, and oil droplets position**

Spindle displacement and rotation time-lapses were rotated to align the initial spindle axis to the horizontal X-axis. Magnet tip position was recorded in DIC and the position of bead aggregates were tracked from their fluorescence signal. Spindle position and orientation were processed manually in Fiji by tracking the centers of two smooth disks, which correspond to spindle poles in the DIC channel. This allowed to compute spindle length, center as well as angles with the magnetic force axis. Spindle displacements and magnetic forces were then projected along the horizontal X-axis for parallel pulling or along the Y-axis for orthogonal pulling. Chromosome plate position was also tracked but we often noticed some small delay in the motion of chromosomes as compared to that of spindle poles, presumably caused by some time-scales associated with internal elastic structures that hold chromosomes. Oil droplets were tracked using the TrackMate plugin in Fiji (6) when the contrast was sufficient, and were checked and corrected manually in some cases. Displacement of the oil droplet was corrected when the egg had displaced during the pulling.

**Tracking of MT distance to the cortex**

Airyscan confocal images stained for MTs and F-actin were projected on a mid-section 2 µm thick, and planar spindles were selected. The horizontal distance between MT ends to the actin cortex was measured, by tracing lines along each MTs to the first border of the F-actin signal using Fiji.

MTs contacting the cortex were also counted from confocal images without F-actin staining in larger z-sections of ~30 µm in thickness and the distance between their ends and the closest surface was measured.

**Segmentation of endomembranes from SBF-SEM images**

SBF-SEM images were processed using Ilastik a machine-learning-based interactive tool (7). The Pixel classifier was trained iteratively with 3 labels (ER; yolk & mitochondria; cytosol) on a cropped image until a satisfying accuracy was reached for endomembrane compartments. Pixel classification maps were then extracted from the full scale dataset.

**Viscoelastic parameter calculation**

Spindle displacement and rotation were fitted with a Jeffreys' model using a custom code written in Matlab (Mathworks) to compute viscoelastic parameters. For the rising phase, the spindle position was fitted using:

$$\frac{d(t)}{F(t)} = \frac{1}{\kappa}\left[1 - e^{-\frac{t}{\tau 1}}\right] + \frac{t}{\kappa\tau 2}\ , \qquad \text{(Equation 1)}$$

where *d* is the displacement along the X-axis or Y-axis (for parallel or orthogonal pulling respectively), and F is the magnetic force. This re-scaling of the displacement by force allows to compensate for small variations in force amplitude during each pull, and implicates that we assume that viscoelastic responses are mostly linear (e.g that the restoring stiffness, κ, and the viscoelastic time-scales, $\tau_1$ and $\tau_2$ are independent of force amplitude). These fits allowed to compute the restoring stiffness, κ, and the viscoelastic time-scales, $\tau_1$ and $\tau_2$ corresponded to the Jeffreys' viscoelastic timescales, allowing to compute viscous drags on spindles as $\gamma = \kappa\tau_1$

Spindle orientation was fitted similarly, using:

$$\frac{\theta(t)}{T(t)} = \frac{1}{\kappa}\left[1 - e^{-\frac{t}{\tau 1}}\right] + \frac{t}{\kappa\tau 2} \qquad \text{(Equation 2)}$$

where θ, is the spindle axis angle and T the torque computed as the force projected orthogonal to the spindle long axis and multiplied by half the spindle length. This allowed to compute a rotational stiffness, κ, and viscoelastic time-scales corresponding to rotational behavior under force.

Spindle relaxation dynamics was fitted using:

$$\frac{d(t)}{d(0)} = (1-a)e^{-\frac{t}{\tau 1}} + a \qquad \text{(Equation 3)}$$

where t = 0 corresponds to the time of the end of force application, to compute the relaxation offset *a* and the decay time-scale $\tau_1$. The same equation was used for rotational relaxation but replacing d by θ.

Fit of the data were obtained for each single force experiment, for spindles or oil droplets, using Nonlinear least squares method in Matlab curve fitting. Various combinations of start points of fit parameters were tried and chosen in a way that maximum R-square and tighter confidence interval were achieved. Curves were also visually inspected to ensure that they passed through more points and were not biased with respect to the data points. Finally, other 2 or 3-element models than Jeffreys' were also tested, but failed to provide better fit of all behaviors than this model.

To estimate the enhanced drag of spindles associated with the confinement by the cell boundaries, we used an effective spindle radius defined as: $R_s = \frac{1}{2}\sqrt{L_s W_s}$, where $L_s$ and $W_s$ the length and the width of the spindle. The enhanced drag of a rigid sphere inside a spherical confinement, is computed as $\gamma_c = K\gamma$ , were K is a wall correction factor and γ=6πηR the drag in an infinite medium (8). K is computed as

$$K = \frac{1-\lambda^5}{1-\frac{9}{4}\lambda+\frac{5}{2}\lambda^3-\frac{9}{4}\lambda^5+\lambda^6} \qquad \text{(Equation 4)}$$

With λ was the ratio between the spindle radius and that of the egg.

**Flow analysis**

The recorded spindle pulling images in DIC were analyzed using the particle image velocimetry PIVlab tool in Matlab (9). The exterior of the egg was masked to be excluded from the analysis. Contrast limited adaptive histogram equalization (CLAHE) and two dimensional Wiener filter with accordingly windows of 20 and 3 pixels widths were applied on the images in the pre-processing steps for denoising. Image sequences were investigated in the Fourier space by three interrogation windows with 64, 32, and 16 pixels widths and 50% overlapped area. The spline method was used for the window deformation and subpixel resolution obtained by two-dimensional Gaussian fits. The distribution of the velocity components of the vectors for each set was visually inspected and restricted to remove the outliers in the post-processing stage. Moreover, two other filters based on the standard deviation and local median of velocity vectors were applied to validate the vector fields. The output vector fields after smoothing were used for the analysis. The vector field was temporally averaged over the pulling or relaxation phase except when tracking imaginary tracers.

For measuring the shear rate in the pulling phase, the vector field on the half side of the egg close to the magnet was spatially averaged along the pulling axis (Fig S6B and S6G). Shear rate was calculated as the ratio of the maximum of the velocity component in the direction of pulling divided by the distance taken for the velocity component to drop to zero. Maximums of the velocity components were obtained by a one dimensional Gaussian fits. Spatial autocorrelation in direction was computed as $c(r) = <\hat{s}(r+r_0).\hat{s}(r_0)>$, where $\hat{s}$ and $r$ were the unit vector in the direction of velocity and distance, respectively, and the results were averaged over initial positions $r_0$. The crossing point of the spatial autocorrelation function with the x axis was considered as the vortex size. Divergence of the vector field was calculated by summation of partial derivatives of its components using central difference. Vorticity was measured as $\omega = \nabla \times v$ where v was the vector field. Vector fields were also used to follow the path of 100 equally spaced tracers along the line perpendicular to the spindle axis by excluding the spindle itself (10). To this aim, the corresponding velocity vector in the selected point was obtained based on bilinear interpolation. The interpolated velocity vector and initial position were used to solve the ordinary differential equation by Euler's method and find the position of the tracer in the next frame. This procedure was continued through all pulling frames.

To compare advection by flows and diffusion, the Peclet number was calculated as Pe=Lu/D with L, u, and D the characteristic length, flow speed, and diffusion coefficient, respectively. Taking the viscosity of the cytoplasm to be 1,000 times that of the water (11), an effective temperature for the active cytoplasm of ~10 times that of room temperature (12), we computed a diffusion coefficient of $\sim 10^{-2}$ $\mu m^2/s$ for a 1 µm radius vesicle. Measured flow speeds in the cytoplasm was in the order of $10^{-2}$ µm/s and eggs of ~100 µm diameter. This leads to the Peclet number $\sim 10^2$ indicating that advection dominates over diffusion for objects of this size.

**Statistical analysis**

All experiments presented in this MS were repeated at least twice and quantified in a number of cells or events detailed in each figure legend. Statistical and correlation analyses were carried out using Prism 6 (GraphPad Software, La Jolla, CA). Statistically significant differences and tests used depended on whether experiments were paired or not and are reported in figure legends.

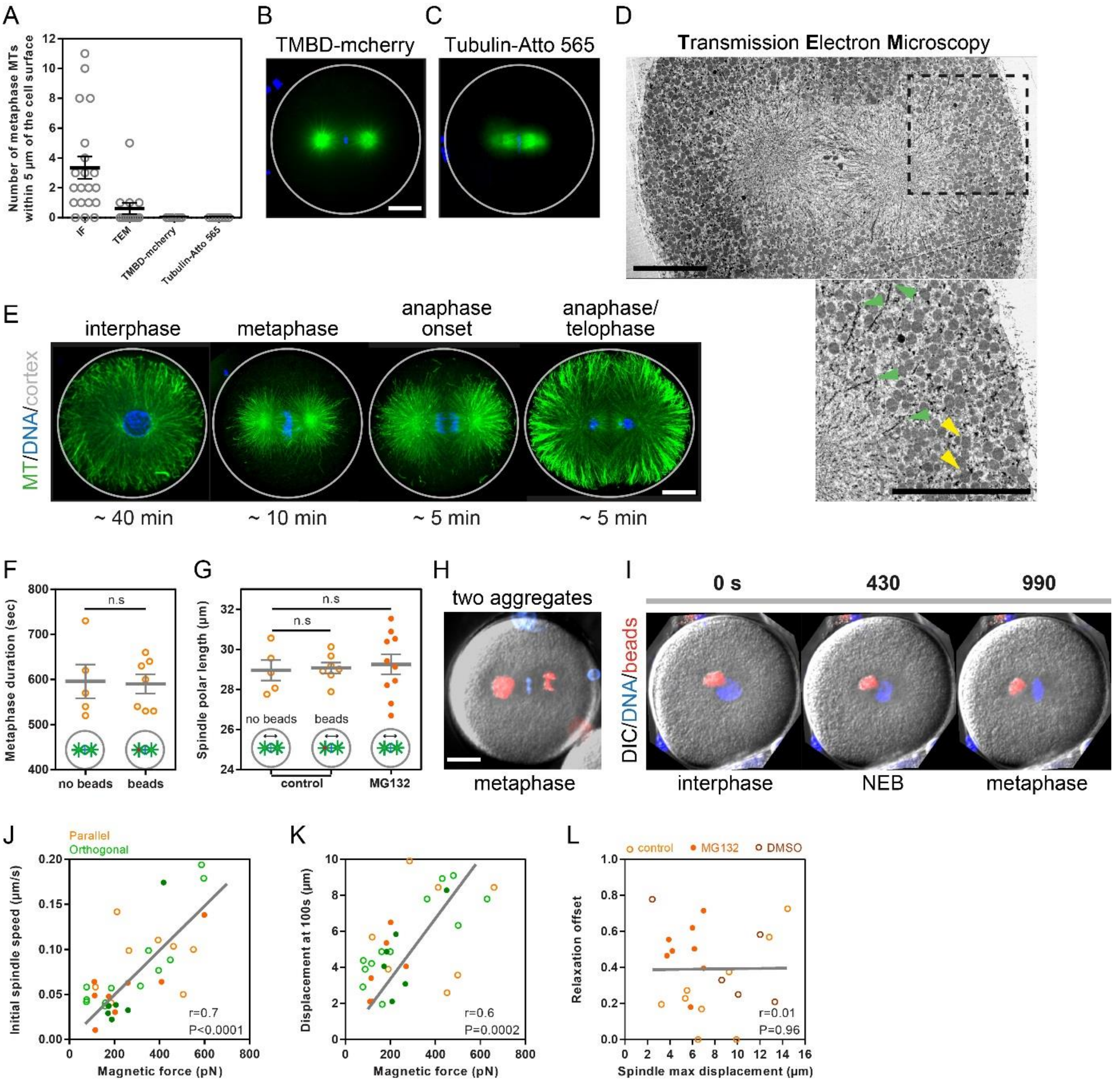

A
Number of metaphase MTs within 5 µm of the cell surface
IF
TEM
TMBD-mcherry
Tubulin-Atto 565
B
TMBD-mcherry
C
Tubulin-Atto 565
D
Transmission Electron Microscopy
E
interphase
metaphase
anaphase onset
anaphase/ telophase
MT/DNA/cortex
~ 40 min
~ 10 min
~ 5 min
~ 5 min
F
Metaphase duration (sec)
n.s
no beads
beads
G
Spindle polar length (µm)
n.s
n.s
no beads
beads
control
MG132
H
two aggregates
metaphase
I
0 s
430
990
DIC/DNA/beads
interphase
NEB
metaphase
J
Parallel
Orthogonal
Initial spindle speed (µm/s)
Magnetic force (pN)
r=0.7
P<0.0001
K
Displacement at 100s (µm)
Magnetic force (pN)
r=0.6
P=0.0002
L
control
MG132
DMSO
Relaxation offset
Spindle max displacement (µm)
r=0.01
P=0.96

**Figure S1. Metaphase astral MTs do not reach the cortex, yet spindles can recoil back to the cell center upon force-induced displacement. (A)** Number of astral MTs reaching a distance less than 5µm to the cortex in metaphase as assayed by Immunofluorescence (IF), Transmission electron microscopy (TEM), or live-imaging of injected TMBD-mCherry or Tublin-Atto-565. **(B-C)** Representative images of metaphase spindles in live cells with MTs labelled by injecting the indicated probes. **(D)** Transmission electron microscopy using fixation methods to reveal the presence of MTs, and close up view with green arrowheads pointing at MT + TIPs, and yellow arrowheads pointing at yolk granules. **(E)** Immunofluorescence of MTs imaged with confocal microscopy in different indicated phases of the cell cycle. The time below the images correspond to the average duration of the cell cycle phases. **(F)** Duration of metaphase in normal zygotes and in zygotes injected with magnetic beads that attach to mitotic spindles. **(G)** Spindle polar length in controls, cells injected with magnetic beads or in cells treated with MG132. **(H)** Example image of an egg in which bead aggregates have split and accumulated on the two spindle poles. **(I)** Time-sequence of a bead aggregate that accumulates at one spindle pole from interphase to metaphase. **(J)** Spindle initial speed under force measured within 30s of force application plotted as a function of the applied force for both orthogonal and parallel pulls (n=35). The line is a linear fit, and correlation coefficients and corresponding P-values are indicated in the plot. **(K)** Displacement after 100s of force application plotted as a function of the applied force for both orthogonal and parallel pulls (n=32). The line is a linear fit, and correlation coefficients and corresponding P-values are indicated in the plot. **(L)** Relaxation offset during spindle recoils is independent of the distance of the spindle to the cortex. The line is a linear fit, and correlation coefficients and corresponding P-values indicated in the plot**.** Error bars correspond to +/- S.E.M. Results were compared by using a two-tailed Mann–Whitney test. n.s, $P > 0.05$.

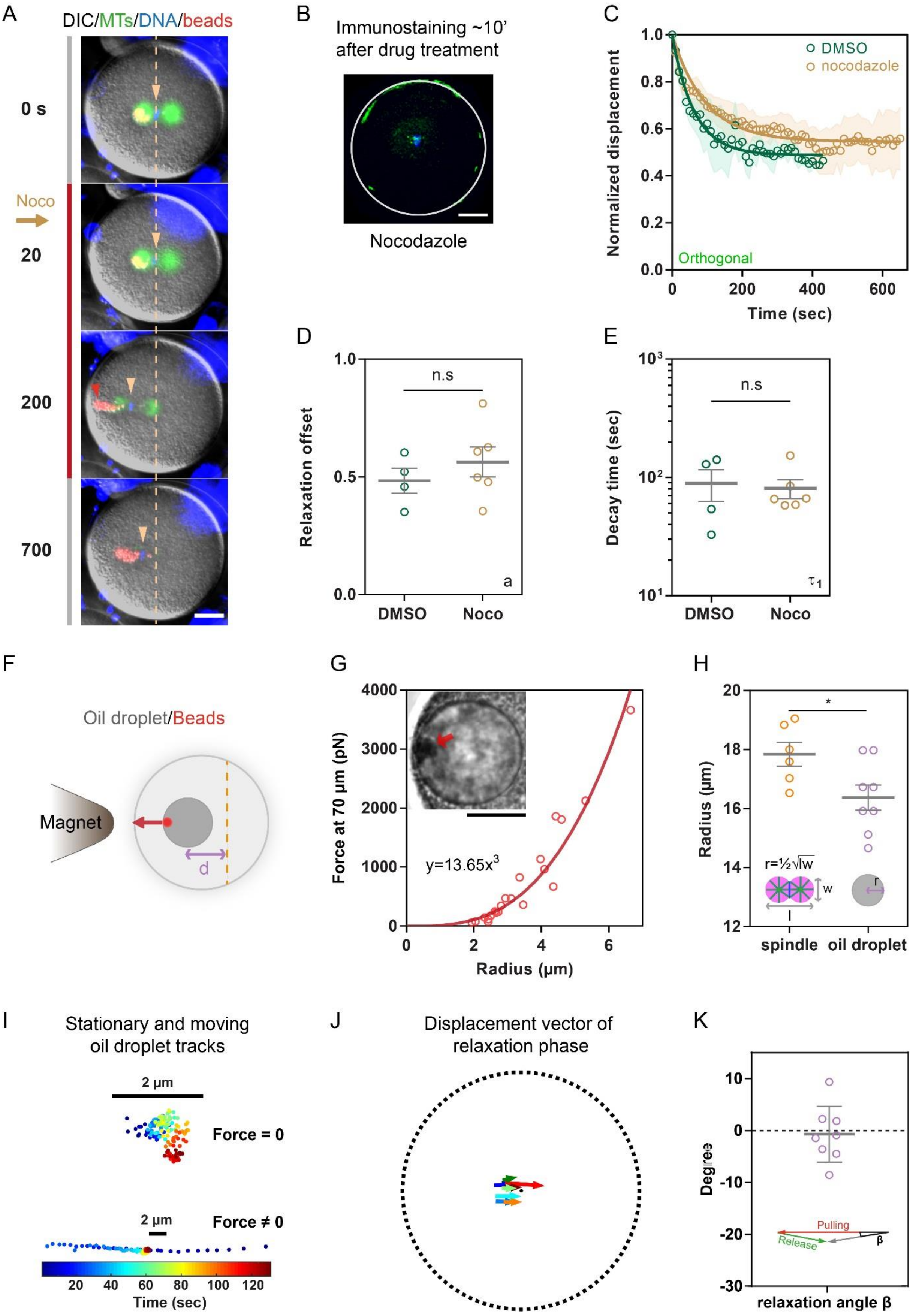

A
DIC/MTs/DNA/beads
0 s
Noco
20
200
700
B
Immunostaining ~10' after drug treatment
Nocodazole
C
Normalized displacement
DMSO
nocodazole
Orthogonal
Time (sec)
D
Relaxation offset
n.s
a
DMSO
Noco
E
Decay time (sec)
n.s
τ1
DMSO
Noco
F
Oil droplet/Beads
Magnet
d
G
Force at 70 μm (pN)
y=13.65x3
Radius (μm)
H
Radius (μm)
*
r=½√lw
w
l
r
spindle
oil droplet
I
Stationary and moving oil droplet tracks
2 μm
Force = 0
2 μm
Force ≠ 0
Time (sec)
J
Displacement vector of relaxation phase
K
Degrees
Pulling
Release
β
relaxation angle β

**Figure S2. Viscoelastic spindle recoils are independent of MT forces, and occur with similar characteristics for a passive oil droplet. (A)** Time-lapse of a spindle pulled with magnetic tweezers and treated during the pull with Nocodazole to depolymerize MTs, and monitor recoils in the absence of MTs. Note the disappearance of MTs. In the experiment the drug was added ~3 min prior to force release to ensure that the effect was maximum at the end of the pulling phase, based on multiple optimization assays. **(B)** Confocal image of a zygote fixed and stained for MTs and DNA after 10' treatment in Nocodazole. **(C)** Time evolution of the normalized displacement back to the cell center in orthogonal pulls when the external force is released in cells treated with the indicated chemicals (n=4 for DMSO and 6 for Nocodazole). **(D-E)** Quantification of the positional offsets to the cell center, and decay time-scales of relaxation curves plotted in C, using a single exponential model. **(F)** Scheme of the assay used to move magnetized oil droplets in cells with magnetic tweezers. **(G)** Close up view of hydrophobic beads aggregates within oil droplets, used to quantify the size of the aggregates, and calibration curve linking the force at 70 µm from the magnet tip to the center of the aggregate. **(H)** Quantification of spindle (n=6) and oil droplets sizes (n=8). **(I)** Tracking of oil droplet centers in the absence and in the presence of external forces. **(J)** Displacement vectors of oil droplets center during the relaxation after force cessation. **(K)** Quantification of the deviation angle between the pulling and relaxation trajectories in individual magnetic oil droplets experiments (n=8). Error bars and shades correspond to +/- S.E.M and +/- S.D/2 respectively. Results were compared by using a two-tailed Mann–Whitney test. n.s, P > 0.05, *, P < 0.05. Scale bars, 20 µm, unless otherwise indicated.

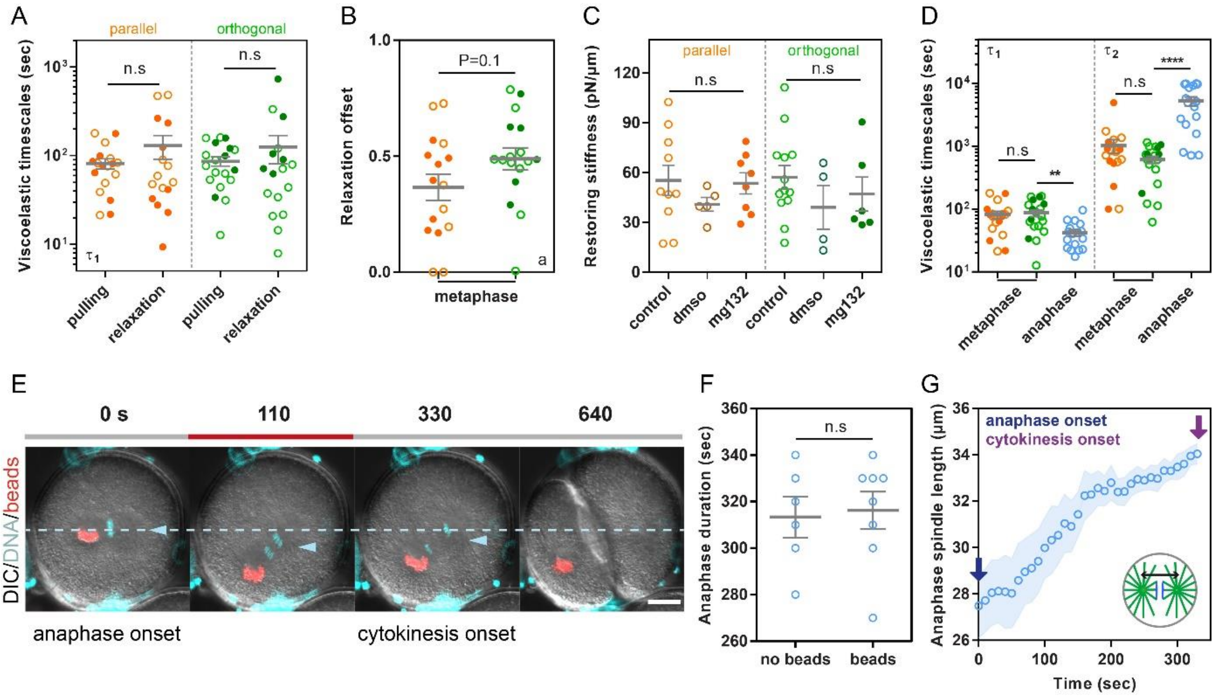

A
parallel
orthogonal
Viscoelastic timescales (sec)
n.s
n.s
τ1
pulling
relaxation
pulling
relaxation
B
Relaxation offset
P=0.1
a
metaphase
C
Restoring stiffness (pN/μm)
parallel
orthogonal
n.s
n.s
control
dmso
mg132
control
dmso
mg132
D
Viscoelastic timescales (sec)
τ1
τ2
n.s
**
n.s
****
metaphase
anaphase
metaphase
anaphase
E
0 s
110
330
640
DIC/DNA/beads
anaphase onset
cytokinesis onset
F
Anaphase duration (sec)
n.s
no beads
beads
G
anaphase onset
cytokinesis onset
Anaphase spindle length (μm)
Time (sec)

**Figure S3. Comparison of metaphase and anaphase spindles pulls and relaxation. (A)** Viscoelastic timescales, $\tau_1$ measured during the pulling or the relaxation phase. **(B)** Relaxation offsets for metaphase spindles pulled parallel or orthogonal. **(C)** Restoring stiffness measured for the indicated conditions in parallel and orthogonal pulls. **(D)** Viscoelastic time-scales for metaphase (n=18 for parallel and 19 for orthogonal pulls) and anaphase spindles pulls (n=17). (**E)** Time-lapse of a representative pulling and relaxing experiment on anaphase spindles. **(F)** Anaphase duration for control eggs vs eggs injected with magnetic beads. **(G)** Time-evolution of anaphase spindle length. Error bars and shades correspond to +/- S.E.M and +/- S.D/2 respectively. Results were compared by using a two-tailed Mann–Whitney test. n.s, P > 0.05, **, P < 0.01, ****, P < 0.0001. Scale bars, 20 µm.

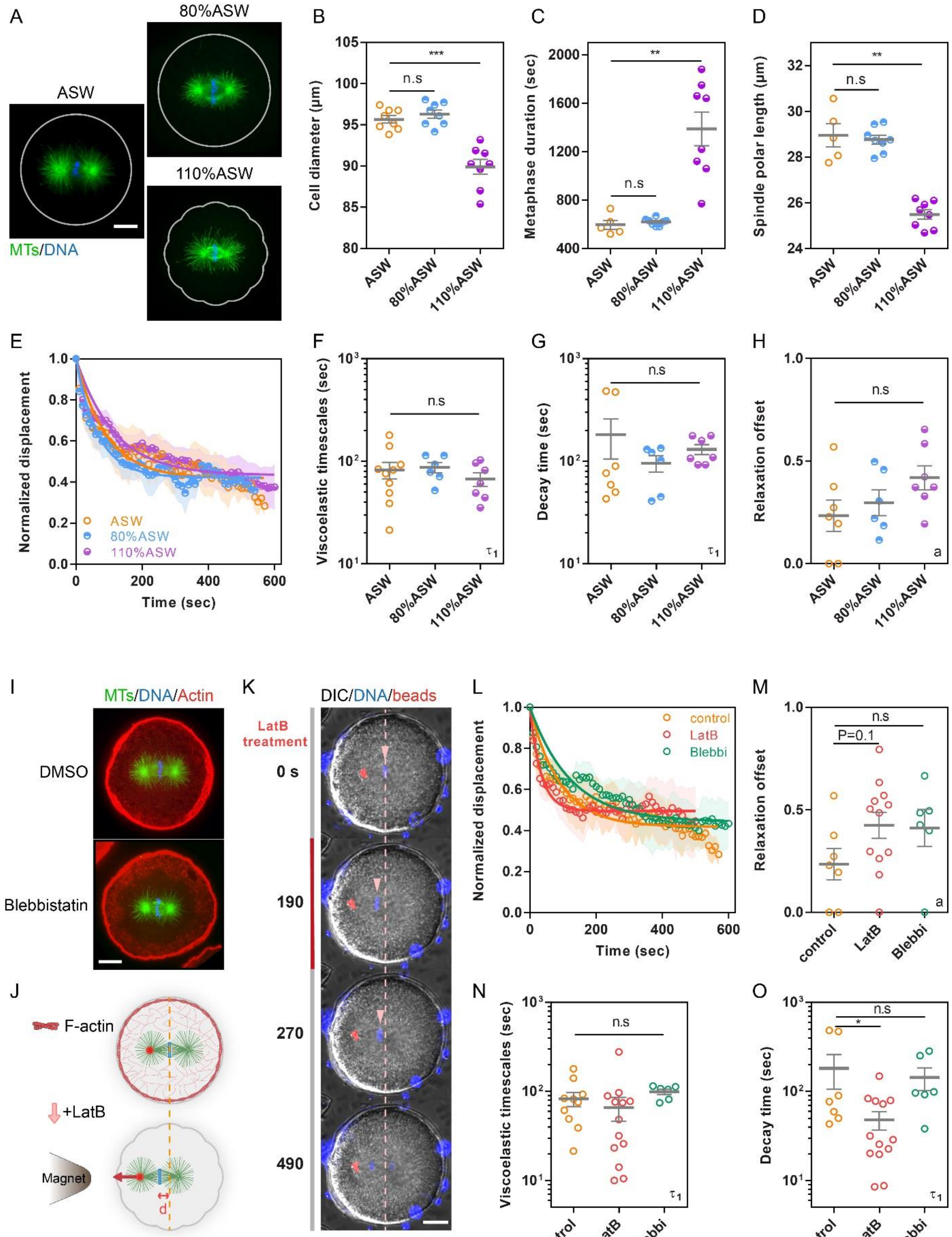

A
ASW
80%ASW
110%ASW
MTs/DNA
B
Cell diameter (μm)
***
n.s
ASW
80%ASW
110%ASW
C
Metaphase duration (sec)
**
n.s
D
Spindle polar length (μm)
**
n.s
E
Normalized displacement
Time (sec)
ASW
80%ASW
110%ASW
F
Viscoelastic timescales (sec)
n.s
$\tau_1$
G
Decay time (sec)
n.s
$\tau_1$
H
Relaxation offset
n.s
a
I
MTs/DNA/Actin
DMSO
Blebbistatin
K
DIC/DNA/beads
LatB treatment
0 s
190
270
490
L
Normalized displacement
Time (sec)
control
LatB
Blebbi
M
Relaxation offset
n.s
P=0.1
a
control
LatB
Blebbi
J
F-actin
+LatB
Magnet
d
N
Viscoelastic timescales (sec)
n.s
$\tau_1$
O
Decay time (sec)
n.s
*
$\tau_1$

**Figure S4. Impact of cytoplasm density, F-actin and myosin II on viscoelastic forces on spindles from bulk cytoplasm. (A)** Confocal images of eggs treated 5min with normal ASW (Artificial Sea Water), 80% ASW or 110%ASW, fixed and stained for MTs and DNA. **(B-D)** Diameter, metaphase duration and spindle length of eggs in ASW (n=8) or 5 min after treatment in 80% ASW (n=8) and 110% ASW (n=8). **(E)** Time evolution of the normalized displacement back to the cell center in controls (n=10) and in cells treated with 80% ASW (n=6) and 110% ASW (n=7). **(F-G)** Viscoelastic timescale, $\tau_1$ measured during the pulling or the relaxation phase, in controls and cells treated with 80% ASW and 110% ASW. **(H)** Relaxation offset for controls and cells treated with 80% ASW and 110% ASW. **(I)** Confocal images of eggs treated with DMSO or Blebbistatin, fixed and stained for MTs, DNA and F-actin. **(J)** Scheme representing F-actin structures and *in vivo* force application on metaphase spindles in the presence of Latrunculin B. **(K)** Time-lapse of eggs treated with Latrunculin B to depolymerize F-actin and assayed for spindle displacement under force and relaxation. **(L)** Time-evolution of the normalized displacement back to the cell center in controls (n=10), in cells treated with Latrunculin B (n=13) and in cells treated with Blebbistatin (n=6). **(M)** Relaxation offset for controls and cells treated with Latrunculin B and Blebbistatin. **(N-O)** Viscoelastic timescale, $\tau_1$ measured during the pulling or the relaxation phase, in controls and cells treated with Latrunculin B and Blebbistatin. Error bars and shades correspond to +/- S.E.M and +/- S.D/2 respectively. Results were compared by using a two-tailed Mann–Whitney test. n.s, P > 0.05, *, P < 0.05, **, P < 0.01, ***, P < 0.001. Scale bars, 20 µm.

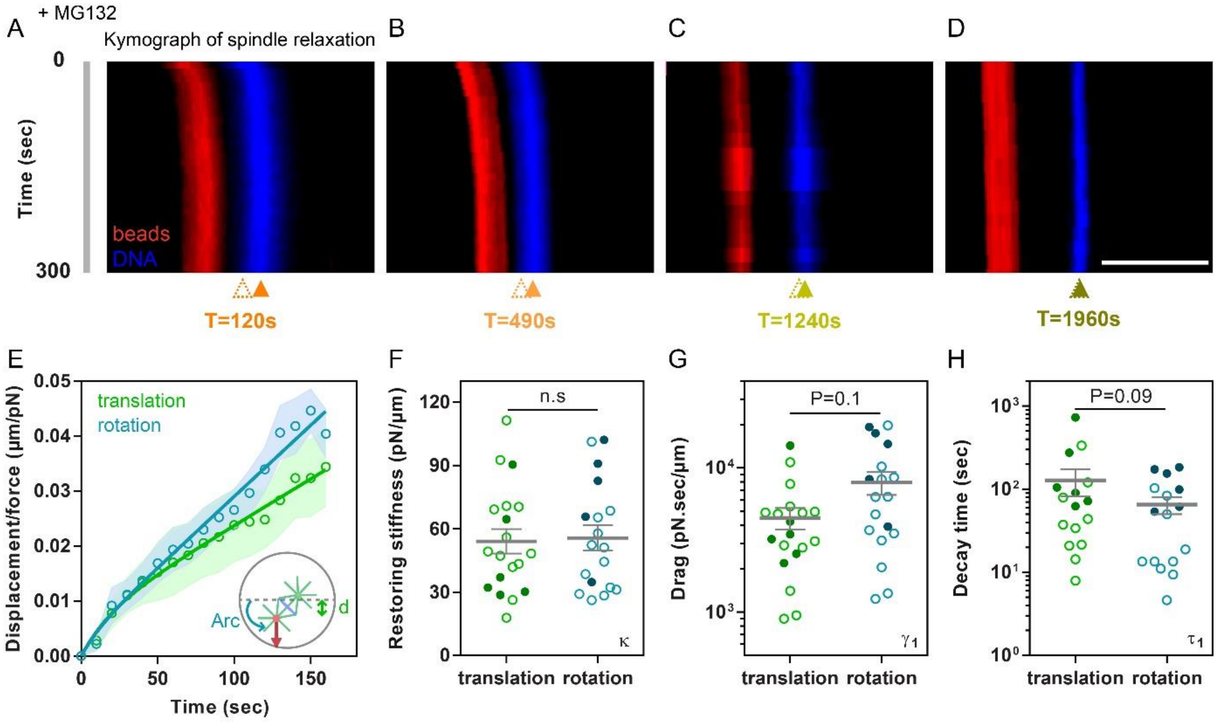

A
+ MG132
Kymograph of spindle relaxation
B
C
D
0
Time (sec)
300
beads
DNA
T=120s
T=490s
T=1240s
T=1960s
E
F
G
H
translation
rotation
Displacement/force (µm/pN)
Time (sec)
Arc
d
Restoring stiffness (pN/µm)
n.s
κ
Drag (pN.sec/µm)
P=0.1
γ1
Decay time (sec)
P=0.09
τ1
translation rotation

**Figure S5. Role of the duration of force application in the dissipation of elastic energy, and differences between translational and rotational behavior. (A-D)** Kymographs following beads and DNA overtime for the relaxation phases upon force applications of varying durations. The dotted arrowhead marks the initial position of spindle centers at the onset of the relaxation phase, and the filled arrowhead indicates the final position after 300s**. (E)** Comparison of translational and rotational creep behavior using the arc length tracking the spindle pole displacement as a measurement of effective displacement during rotational motion. **(F-H)** Restoring stiffness, drags and decay times measured for translation vs rotation. Open and closed dots correspond respectively to data obtained in control or MG132 treated cells. Error bars and shades correspond to +/- S.E.M and +/- S.D/2 respectively. Results were compared by using a two-tailed Wilcoxon test. n.s, $P > 0.05$. Scale bars, 20 µm.

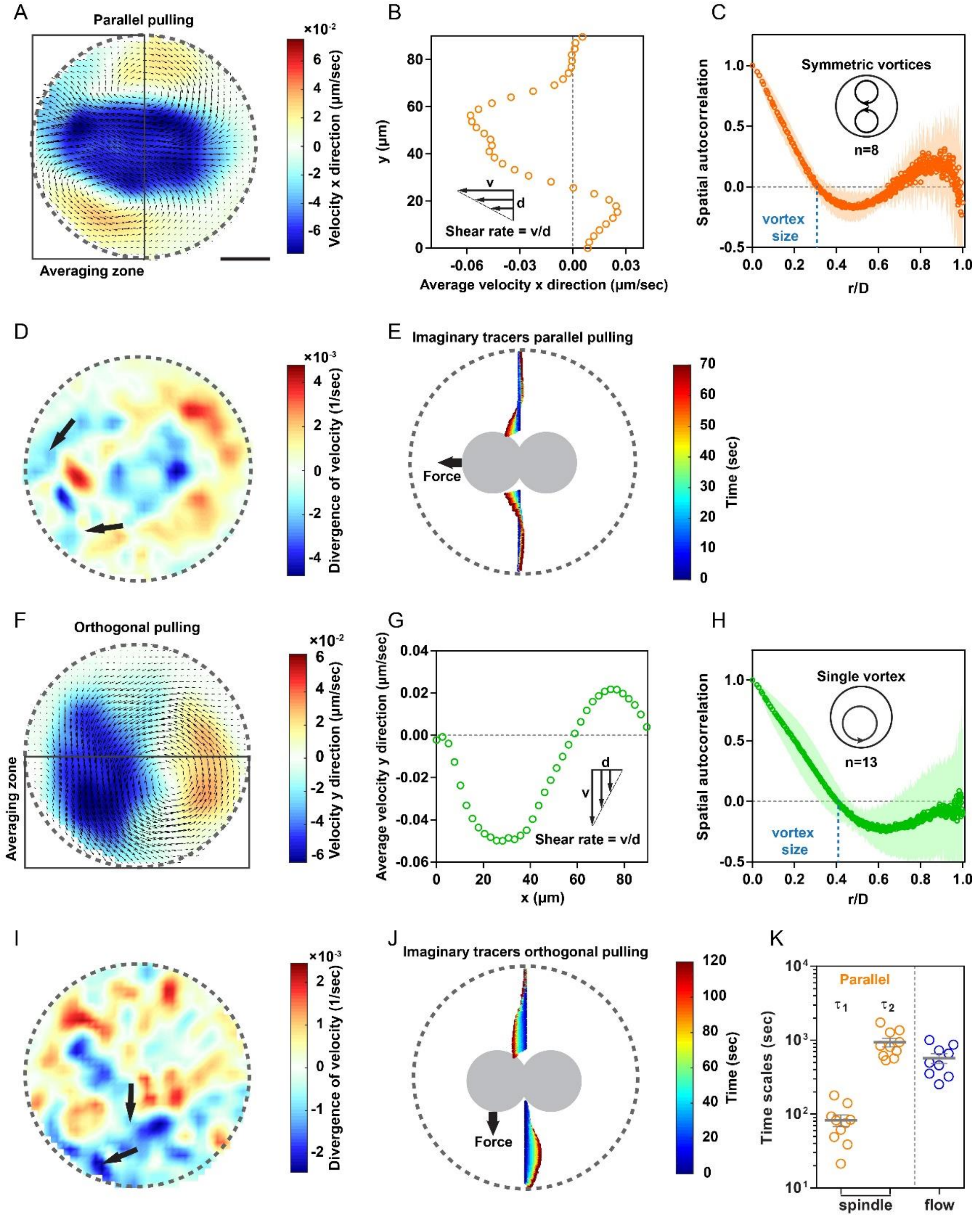
A
Parallel pulling
Averaging zone
×10⁻²
Velocity x direction (µm/sec)
B
y (µm)
Shear rate = v/d
Average velocity x direction (µm/sec)
C
Symmetric vortices
n=8
Spatial autocorrelation
vortex size
r/D
D
×10⁻³
Divergence of velocity (1/sec)
E
Imaginary tracers parallel pulling
Force
Time (sec)
F
Orthogonal pulling
Averaging zone
×10⁻²
Velocity y direction (µm/sec)
G
Average velocity y direction (µm/sec)
Shear rate = v/d
x (µm)
H
Single vortex
n=13
Spatial autocorrelation
vortex size
r/D
I
×10⁻³
Divergence of velocity (1/sec)
J
Imaginary tracers orthogonal pulling
Force
Time (sec)
K
Parallel
τ1
τ2
Time scales (sec)
spindle
flow

**Figure S6. Flows of cytoplasm elements and associated analysis during spindle translational and rotational motions. (A**) Flow maps of a representative spindle pulled with magnetic forces parallel to its axis, superimposed with a color map of the flow velocity projected along the x-axis. **(B)** Averaged velocity component in the horizontal direction (x) in the box indicated in A, plotted as a function of the vertical distance (y), used to compute shear rates. **(C)** Spatial autocorrelation averaged on 8 individual experiments and flow analysis used to compute flow vortex sizes. **(D)** Color maps of the divergence of the velocity used to locally map zones where the cytoplasm matrix may compress (indicated with arrows). **(E)** Trajectories of imaginary flow tracers placed along the direction orthogonal to the spindles used to visualize net displacement of cytoplasm material as a function to the distance to the spindle. **(F-J)** Similar analysis as in (A-E) for spindles pulled orthogonal to the spindle axis. **(K)** Viscoelastic timescales $\tau_1$ and $\tau_2$ extracted for spindles from rising curves and compared to a time-scale calculated from the inverse of the shear rate from the flow maps. Error bars and shades correspond to +/- S.E.M and +/- S.D respectively. Scale bars, 20 µm.

**Supporting movie legends**

**Movie S1. Metaphase spindles remain still in the cell center.** Time-lapse of a zygote with DNA labelled with Hoechst, through metaphase, followed by a time-lapse of an egg treated with MG132 to prolong metaphase. Time is in min:sec.

**Movie S2. Metaphase spindle pulled parallel to their long axis with magnetic tweezers in live cells.** Time-lapse of 2 metaphase spindles pulled with magnetic forces along their long axis and let to recoil and divide. Beads are labelled in red and DNA in blue. Time is in min:sec.

**Movie S3. Metaphase spindle pulled orthogonal to their long axis with magnetic tweezers in live cells.** Time-lapse of 2 metaphase spindles pulled with magnetic forces orthogonal to their long axis, let to recoil and divide. Beads are labelled in red and DNA in blue. Time is in min:sec.

**Movie S4. Magnetized oil droplets pulled in the cytoplasm and let to recoil.** Time-lapse of an egg injected with an oil droplet containing magnetic beads, pulled with magnetic tweezers and let to recoil. Time is in min:sec.

**Movie S5. Serial Block Face images of metaphase cytoplasmic components with corresponding 3D pixel classification.** Z-stacks of the metaphase cytoplasm with classified chromosomes (blue), ER (purple), Yolk (yellow) and Mitochondria (red). Close up views of the spindle region are presented at bottom.

**Movie S6. Cytoplasm flows during parallel and orthogonal pulls.** Time-lapse of an egg in which the metaphase spindle was pulled parallel to its long-axis, superimposed by the flow vector maps in red; followed by a similar time-lapse for an orthogonal pull. Time is in min:sec.
